# *mergem*: merging and comparing genome-scale metabolic models using universal identifiers

**DOI:** 10.1101/2022.07.14.499633

**Authors:** Archana Hari, Daniel Lobo

## Abstract

Numerous methods exist to produce and refine genome-scale metabolic models. However, due to the use of incompatible identifier systems for metabolites and reactions, computing and visualizing the metabolic differences and similarities of such models is a current challenge. Furthermore, there is a lack of automated tools that can combine the strengths of multiple reconstruction pipelines into a curated single comprehensive model by merging different drafts, which possibly use incompatible namespaces. Here we present *mergem*, a novel method to compare and merge two or more metabolic models. Using a universal metabolic identifier mapping system constructed from multiple metabolic databases, *mergem* robustly can compare models from different pipelines and merge their common elements. *mergem* is implemented as a command line tool, a Python package, and on the web-application Fluxer, which allows simulating and visually comparing multiple models with different interactive flux graphs. The ability to merge and compare diverse genome scale metabolic models can facilitate the curation of comprehensive reconstructions and the discovery of unique and common metabolic features among different organisms.

## 1. Background

Genome scale metabolic models (GEMs) are *in silico* descriptions that can represent and simulate the metabolic networks of biological systems at the cellular or even organismal level. Each model comprises a set of mathematically formulated gene-protein-reaction relationships that contribute to the metabolic state of the biological system (1, 2). There are four major stages for reconstructing GEMs: draft reconstruction, conversion to mathematical format, refinement, and network evaluation. The last three steps are typically iterated until predictions match experimental findings and the validated model is considered a robust GEM (3). All the metabolic information available of a particular system should be encapsulated within robust GEMs, with the goal for the models to be metabolically complete (4). Several pipelines and approaches to generate draft reconstructions are available, including fully automatic tools (5). However, reconstructions built from the same genome using different tools frequently result in different metabolic coverage due to differences in the underlying algorithms and reaction database (6). Merging such draft reconstructions for the same organism into a single comprehensive model can increase the metabolic coverage, leverage the advantages of different reconstruction tools, and thereby improve the completeness of the resulting model. The Human Metabolic Atlas (7), an integration of HMR2, iHSA, and Recon3D GEMs, is testament to the need for merging and comparing multiple models. Yet, the large size and connectivity of GEMs and the use of different identifier (ID) namespaces—for reactions, metabolites, and genes—by different pipelines makes the merging and visualization of different reconstructed GEMs a current challenge (8, 9).

A number of tools are available that can merge GEMs and metabolic reconstructions, but their functionality is limited. MetaMerge (10) is a Python library that can take two metabolic networks as input and unify their metabolites and reactions based on their features such as metabolite CAS number, KEGG ID, IUPAC name, reaction name, gene name, or pathway name. modelBorgifier (11) is a MATLAB toolbox for matching and comparing, semi-automatically, two given genome-scale reconstructions by merging metabolites and reactions based on a set of parameters such as metabolite name, KEGG ID, reaction name and the number of reactants and products in reaction. iMet (12) is a graphical user interface software that merges metabolic networks in a semi-automatic fashion. The MetaNetX web-interface (13) provides tools for combining models and identifying common and unique components between pairs of models. The COBRApy (14) package also offers a merge function, but it is based on merging reactions and metabolites with the same IDs. Although these tools are very useful for processing GEMs, they have important limitations that reduce their applicability (see section 2.4 for a detailed comparison). In brief, all these tools can merge only two models at a time, have problems merging models with different database IDs, can take a long time to merge two models, do not offer a visual comparison, and most of them require the user to be familiar with a programming language such as MATLAB or Python.

The comparison and merging of models can be aided by visualizing the commonalities and differences of metabolite and reaction components in each model. Such graphical representations can be beneficial not only for unifying draft reconstructions from different pipelines, but also to discover differences between cells and organisms with validated GEMs. While tools such as Escher (15), MetExplore (16), and Pathview (17) can be used to visualize GEMs, they are not amenable for comparing the differences between multiple models or merging them. There is thus a need for a new approach that can robustly compare, merge, and visualize multiple reconstructions produced from different pipelines with different database IDs and without keeping duplicates or causing loss of information.

Here we present a novel methodology and software tool called *mergem* for comparing and merging GEMs and draft reconstructions. The proposed algorithm translates and unifies metabolite IDs using a comprehensive database mapping system and then compares reactions based on their participating reactant and product metabolites. The method is freely available as a Python package and command line tool. In addition, *mergem* has been integrated into the user-friendly web application Fluxer (18), which allows any user without programming knowledge to perform the merging, comparison, and visualization of any number of models directly in the web application. Figure 1 illustrates the main steps of the proposed methodology. The results section demonstrates the application of the method with several use-cases, including merging multiple draft reconstructions using different namespaces and comparing published GEMs from different organisms. The proposed merging algorithm and tool and its integration with Fluxer’s robust metabolic network visualization and analysis capabilities can not only streamline the reconstruction of robust GEMs but also visually compare flux networks of different reconstructions and organisms with a user-friendly graphical interface. In this way, *mergem* can facilitate the generation of comprehensive reconstructions and novel hypotheses for metabolic engineering, biomedicine, and drug development applications.

**Figure 1.**
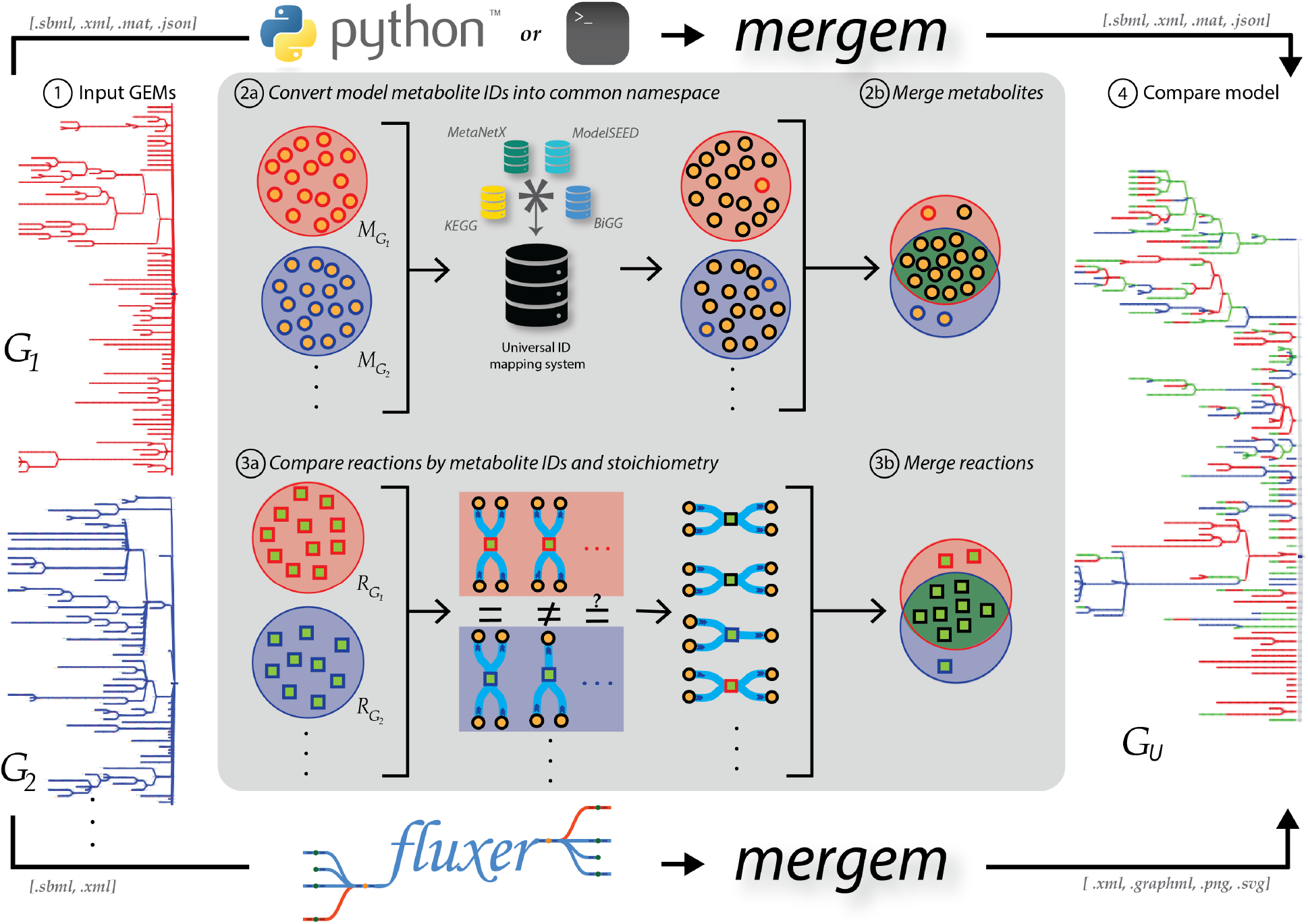
Key steps of the mergem method for merging genome-scale metabolic models (GEMs), as implemented in a Python package, a command-line tool, and Fluxer user-friendly web application. **1.** Two or more GEMs are taken as input. **2a.** The model metabolite IDs are mapped to a universal common namespace. **2b.** Metabolites with the same universal ID are merged. **3a.** Reactions are compared using reactant and product metabolite universal IDs. **3b.** Similar reactions are merged. **4.** The resultant merged model is returned, along with merging information and a Jaccard matrix measuring the distance between each pair of input models.

## 2. Methods

### 2.1. Jaccard Distance between GEMs

The differences between two GEMs in terms of common metabolites and reactions can be quantified using the Jaccard distance (19) as

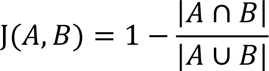

where, *A* and *B* represent either the metabolite or reaction sets for each of the models. The Jaccard distance range is [0,1], where 0 indicates that the sets (metabolites or reactions) are identical and 1 indicates that none of the elements are matched between the models.

Since the Jaccard distance is commutative and equal to zero when applied to the same set, we define a Jaccard distance matrix JM for a set of *n* GEMs *G*_1_,…, *G*_n_ as

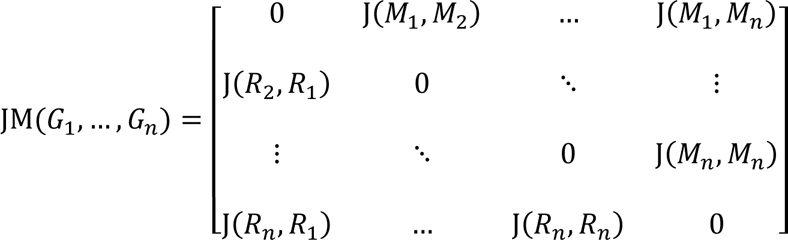

where *M*_i_ and *R*_i_ denote the set of metabolites and reactions, respectively, in the model *G*_i_.

### 2.2. Algorithm for merging models

The *mergem* algorithm for merging two or more GEMs can be summarized into five major steps:

1. Template model assignment
2. Merging subsequent input models
3. Assigning objective function to merged model
4. Calculating Jaccard distances between pairs of models
5. Returning merged model, metabolite and reaction mappings, and Jaccard distance matrix

#### 2.2.1. Definitions

##### GEM

A GEM *G* is a system (*M*_G_, *R*_G_, *M*_G_), where *M*_G_ is a set of metabolites, *R*_G_ is a set of metabolic reactions, and *M*_G_ is a set of objective reactions in *G*. A reaction *r* is an ordered pair (*T*_r_, *P*_r_), where *T*_r_ is a set of reactant metabolites and *P*_r_ is a set of product metabolites. *S*(*m*, *T*_r_) and *S*(*m*, *P*_r_) denote the stoichiometry of metabolite *m* in reactants *T*_r_ or products *P*_r_, respectively.

##### mergem ID

A *mergem* ID is a string formed with the concatenation of a universal ID (from the mapping system derived in Section 3.1) and the subsystem that the molecule is found in (*e.g.*, cytoplasm, extracellular, mitochondria). For example, a hydrogen atom in the cytoplasm is represented as *mergem_78_c*. If no universal ID exists for the metabolite, its original ID is used instead. Formally, ID(m) denotes the ID of metabolite *m* and ID_0_(*m*) denotes its mapped *mergem* ID. This naming system is only internal to the algorithm for merging common metabolites from the input models to overcome the problem with models using different namespaces. The output merged model does not contain any *mergem* ID, since the *mergem* IDs are reverted to the original ID used by the first input model containing it. The original metabolite IDs are temporarily stored for reversion as *σ*: *m* → ID(m).

##### Reaction key

A reaction key contains the *mergem* IDs of its participating metabolites, each paired with an integer denoting it as a reactant (−1) or product (1) in the reaction. In particular, a reaction key K(*r*) of a reaction *r* is a set of pairs {(*ID*_θ_(*m*_i_), −1),… (*ID*_θ_(*m*_j_), 1),… }|*m*_i_ ∈ *T*_r_, *m*_j_ ∈ *P*_r_. For example, the key for the reaction

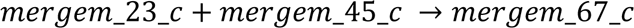

would be

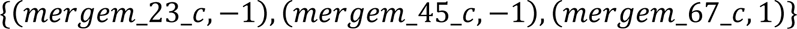

Reaction databases such as MetaNetX explicitly include hydrogen atoms in reaction products and reactants, but others do not include them. Further, MetaNetX models contain a special model compartment called ‘boundary’ to represent model boundary that is independent of other commonly found biological compartments in GEMs, such as ‘cytoplasm’, ‘mitochondria’, or ‘extracellular’. Since these conventions are not standardized across all reaction databases, the *mergem* algorithm is designed to ignore explicit hydrogen atoms and boundary metabolites when generating reaction keys, which allows for better reaction matching across databases.

##### Source mapping dictionaries

For analyzing the differences and commonalities between the input models, the algorithm tracks the presence of each merged metabolite and reaction in each input model. These mappings are formally defined for metabolites with _μ_: *m* ∈ *M*_∪_ → G^∗^ ⊆ {*G*_1_,…, *G*_n_} and for reactions with _ρ_: *r* ∈ *R*_∪_ → G^∗^ ⊆ {*G*_1_,…, *G*_n_}.

##### Algorithm input

A tuple of *n* GEMs [*G*_1_,…, *G*_n_] to merge and whether the objective should be merged or selected.

##### Algorithm output

The merged GEM *G*_∪_, a source mapping of metabolites in the merged model to their original models _μ_: *m* ∈ *M*_∪_ → G^∗^ ⊆ {*G*_1_,…, *G*_n_}, a source mapping of reactions in the merged model to their original models _ρ_: *r* ∈ *R*_∪_ → G^∗^ ⊆ {*G*_1_,…, *G*_n_}, and a Jaccard distance matrix *JM* with all pair-wise metabolite and reaction distances between the input models.

#### 2.2.2. Assigning and translating the template model

The *mergem* algorithm begins by initializing the first input model as the template for merging.

##### Template metabolites

Each metabolite in the template model is translated into its corresponding *mergem* ID and copied to the merged model. Each translation is performed only after checking that a metabolite with the same *mergem* ID is not already present in the merged model. This can happen when the model contains different metabolites that map to the same *mergem* ID. For example, NH2 and NH3 map to the same *mergem* ID since many databases consider them synonymous due to their difference in a single proton. In these cases, the original metabolite ID is retained and not translated. This guarantees that the merged model contains all the metabolites from the template (first) model and that the rest of the models can merge the same conflicting metabolites to the template model correctly.

##### Template reactions

All the template model reactions are copied to the merged model, and their reaction keys generated for comparison with subsequent models. Notice that reaction keys always contain the translated *mergem* IDs, independently if different metabolites are translated to the same *mergem* ID. This maximizes the ability of the algorithm to merge equivalent reactions.

#### 2.2.3. Merging models

Subsequent input models are then merged into the merged model by translating their metabolites to *mergem* IDs and generating keys for their reactions.

##### Metabolite translation

Metabolites in the non-template models are translated to *mergem* IDs, except when their original IDs are not found in the universal ID mapping system or they already exist in the merged model— which can occur when multiple metabolites in the template model map to the same *mergem* ID. In these cases, their original ID is maintained according to the priority established by the order of the input models. Similar to the template model, different metabolites in a model to merge can map to the same *mergem* ID. In this case, the reactions in the model to merge are updated with the metabolite labeled with the new *mergem* ID. An exception to this rule is when the same reaction contains both metabolites and such substitution would affect the stoichiometry (or even remove the metabolite altogether from the reaction if their stoichiometries cancel out). In this case, the metabolite original ID is maintained. This methodology results in maximizing correct reaction matchings while conserving original reactions when conflicts appear in translated metabolite IDs.

##### Reaction matching and merging

Reactions not in the model objective function (which are processed separately) are merged by adding new reactions to the merged model only when their reaction key does not exist among the reactions already included in the merged model. Both the forward and reverse keys are considered a match since their bounds define the actual flux direction in a model. Reaction IDs are not changed during the merging process, except when another reaction with the same ID already exists in the merged model. In this case, the symbol ‘∼’ is appended to the ID of the new reaction to avoid an ID conflict.

#### 2.2.4. Assigning an objective function

After all the input models have been merged, an objective function is added to the merged model according to the criteria selected by the user: either a single objective function copied from one of the input models or a merged objective function that is the average from all the input models.

##### Single objective function

If the user selected an objective function from one of the input models, all the objective reactions from that model are added to the merged model and the merged model is assigned the same objective function.

##### Merged objective function

If the user selected to merge all objective functions, all the reactions in the objective functions from all the input models are merged into a single reaction. Metabolites included in multiple reactions are included only once in the merged objective reaction and its stoichiometry is averaged among all the objective reactions containing it. This option is particularly useful for merging and comparing biomass functions and compositions between different models. A merged model with a merged biomass objective function includes an average proportion of precursors among the input models for cells to grow, which can then be used to compute reaction fluxes.

### 2.3. Algorithm pseudocode

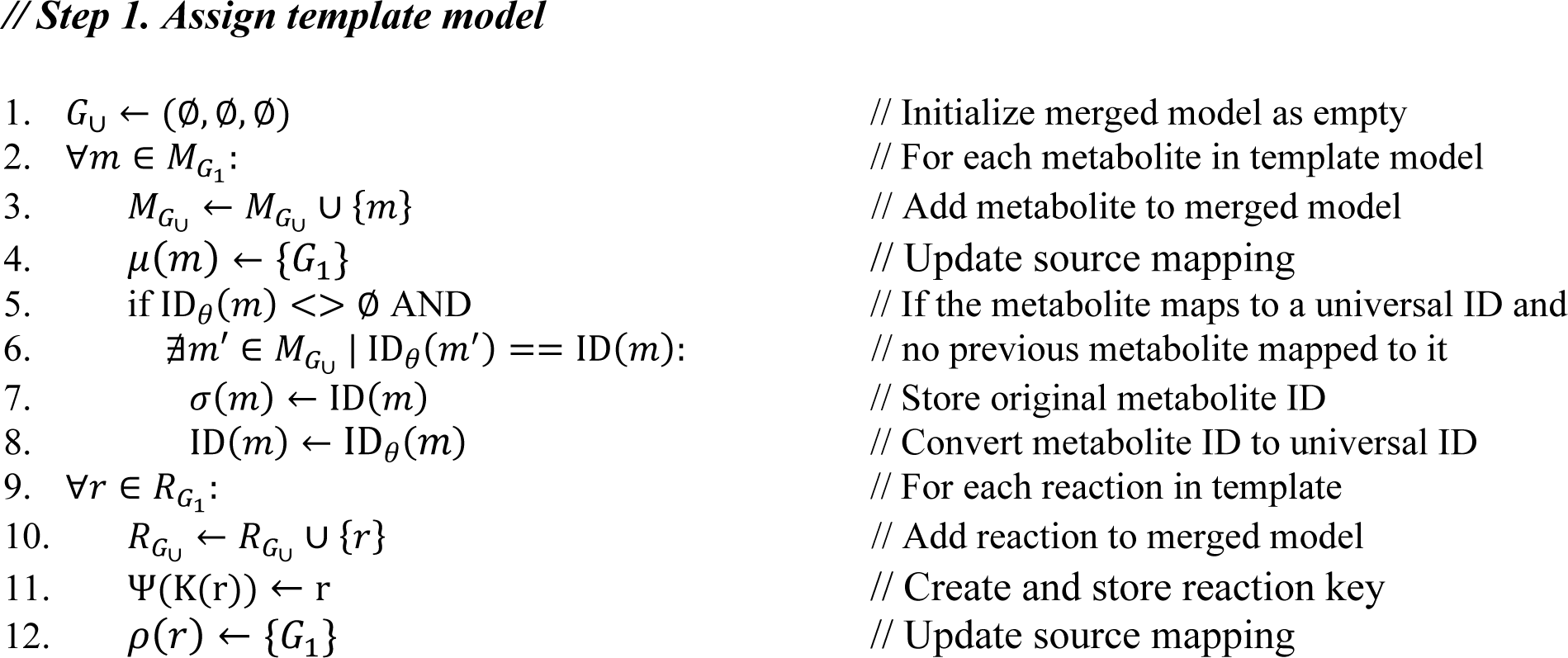
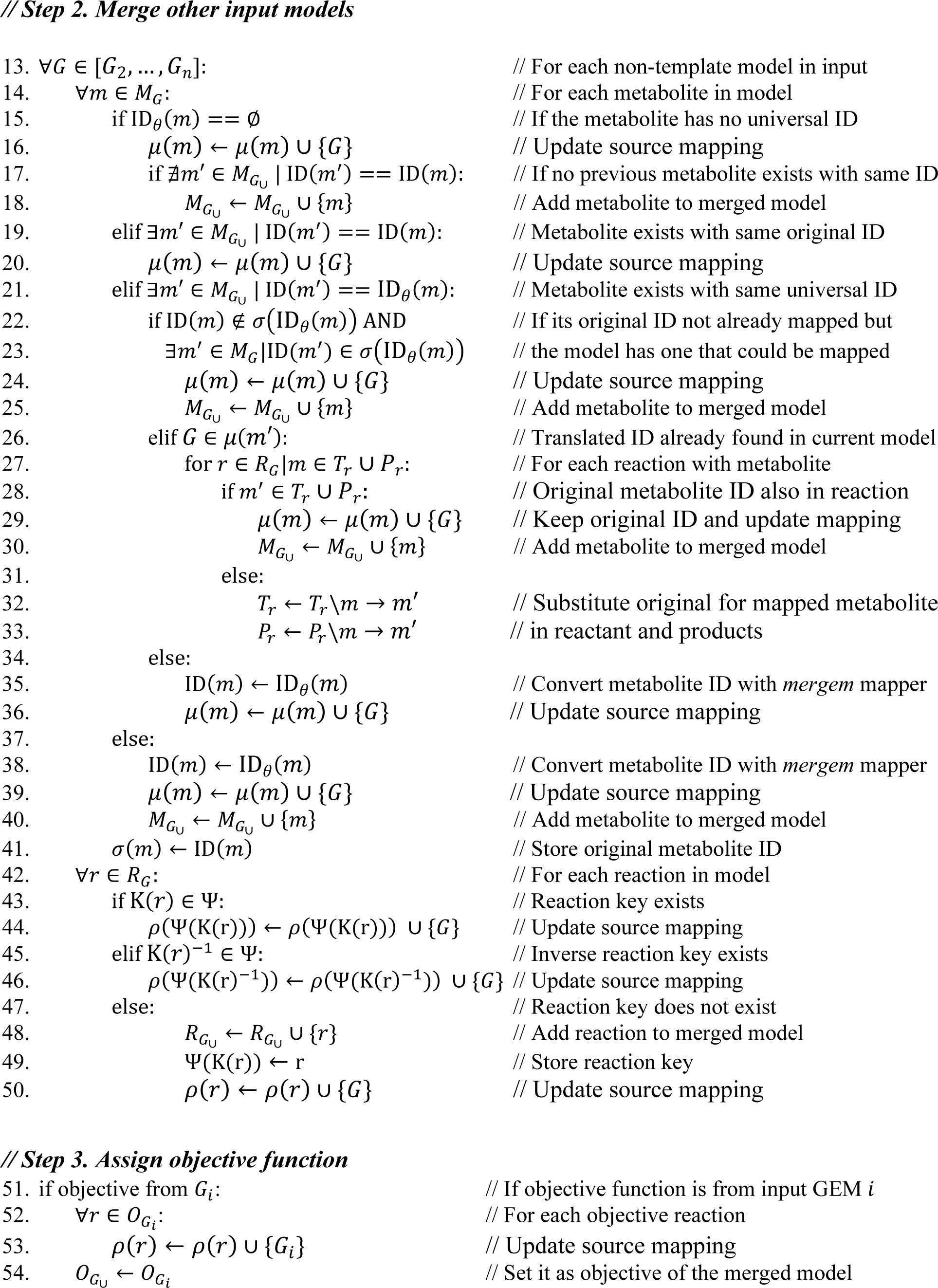
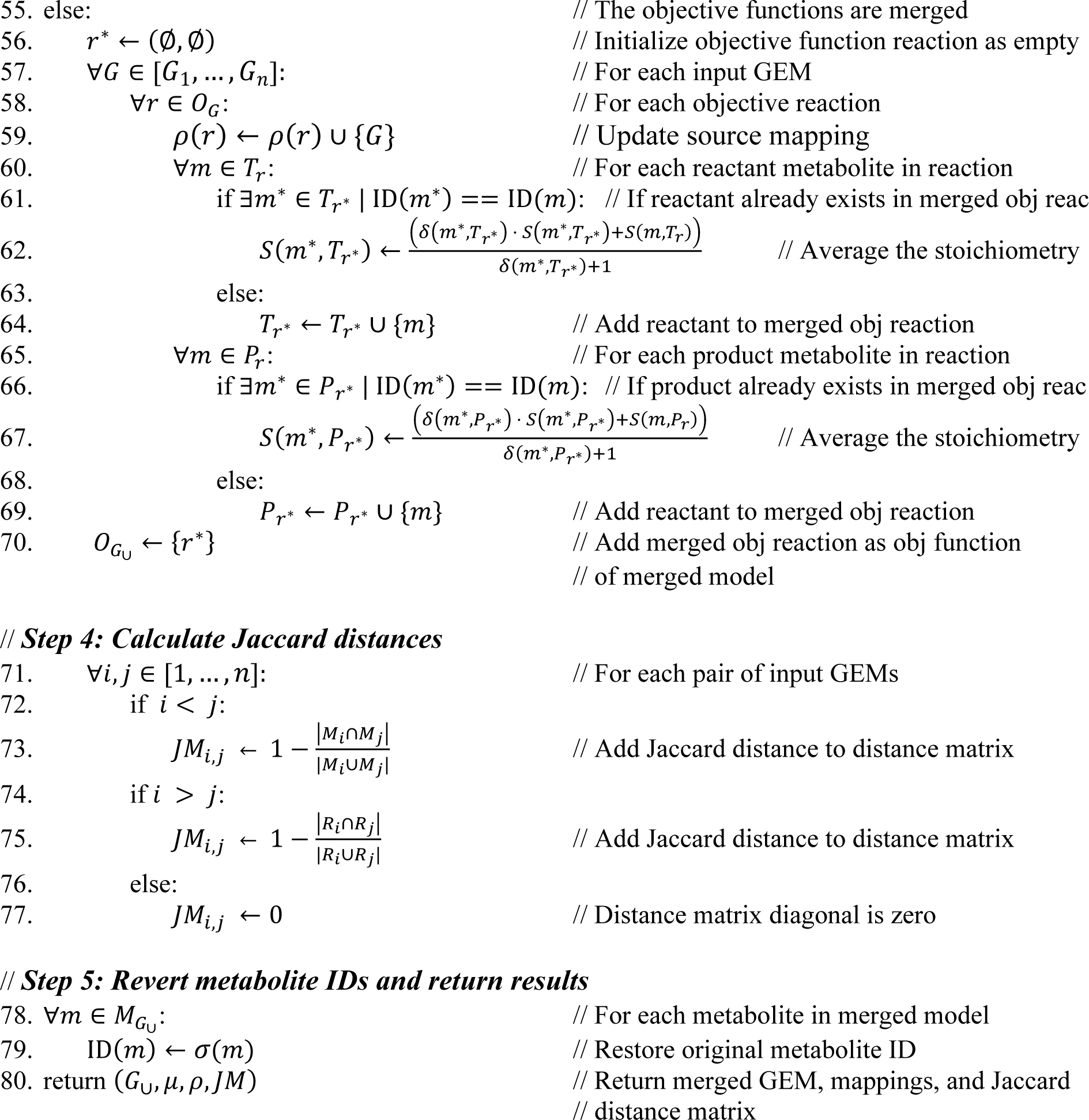

### 2.4. *mergem* Python package

The merging and comparison algorithm has been implemented as the open-source Python package *mergem*, which is freely available in the Python Package Index, PyPI (https://pypi.org/project/mergem/). The package can be executed on the command-line or imported into Python scripts. Depending on the execution method, users can provide a list of COBRApy (20) model objects or filenames (in SBML, MATLAB, or JSON formats) as input to *mergem* along with the objective function to use. The command-line execution automatically saves the resulting merged model as an SBML file, or any other output filename and format specified by the user for maximum compatibility with other tools. The package also computes and returns the Jaccard distance matrix between the models and dictionaries with the source models for each metabolite and reaction, as well as provides methods to translate metabolite and reaction universal IDs and retrieve their consolidated properties.

### 2.5. *mergem* in Fluxer web interface

*mergem* has been integrated in the web-application Fluxer, enabling merging input metabolic models and identifying unique and common metabolic components within the input models as well as for retrieving their properties. Fluxer (18) uses HTML5 and JavaScript for the front end and Python and Flask (Pallets) for the backend. Models loaded into the application are stored in an internal SQLite database. The COBRApy (20) package is used to read, FBA-optimize (21), and write the SBML model files. Graph layouts are visualized with the D3.js library (22). FROG reports are generated with the *fbc_curation* package (23).

## 3. Results

### 3.1. Universal mapping of metabolite and reaction IDs

Different draft reconstruction pipelines label metabolites and reactions with different sets of IDs, which preclude their direct use for merging and comparing metabolites and reactions. Various efforts including ModelSEED biochemistry (24), MetaNetX/MNXref (25, 26) and BiGG (27) exist to overcome the ID inconsistency problem by including mappings from their native IDs with those from other resources. However, there is no streamlined method to apply such mappings from different databases, which also contain inconsistencies (8, 9).

To overcome the problems with multiple ID systems and inconsistent mappings, we developed an algorithm and implemented it in *mergem* to automatically process and reconcile database IDs, resulting in a universal mapper for metabolites and reactions. The tool downloads the most recent data from the main databases for metabolic information, including MetaNetX, ModelSEED, BiGG, KEGG (28), and ChEBI (29), and iteratively processes their metabolite and reaction information, as well as cross-reference mappings. Metabolite and reaction entities are extracted from each database and assigned a unique ID (called universal ID) linked to their listed set of properties (name, molecular weight, InChI key (30), chemical formula, and E.C. number). Table 1 shows the specific metabolite and reaction properties obtained by *mergem* from each database.

**Table 1.**
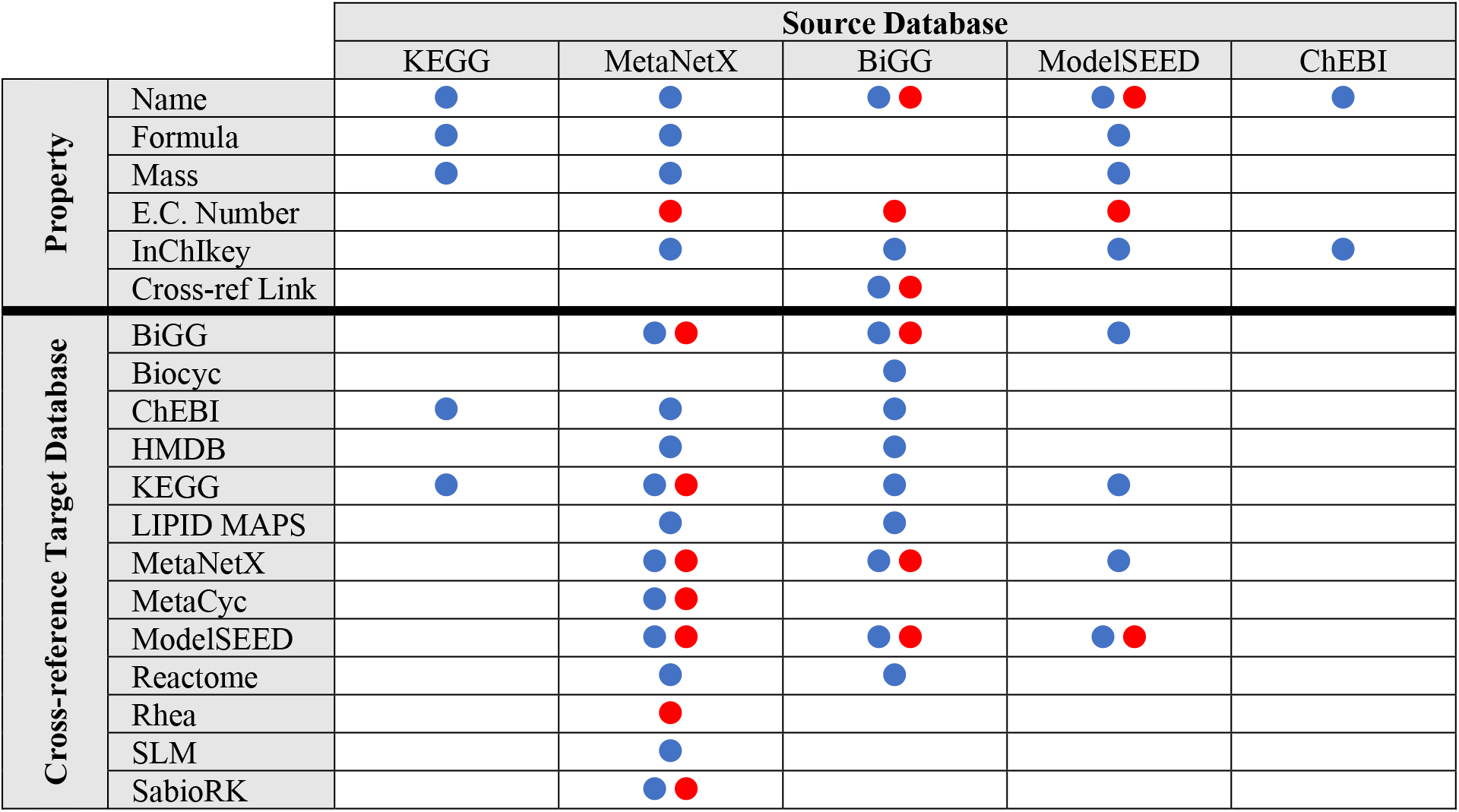
mergem automatically downloads, processes, and consolidates metabolite (blue dot) and reaction (red dot) properties and IDs from source databases into a universal ID mapper for merging models.

Next, the algorithm processes the cross-reference mappings listed in each database as pairs of metabolite or reaction IDs linking one native database ID (source) with an ID from another database (target). Table 1 shows the source and target databases used for ID mappings. Secondary IDs and mappings from ModelSEED are not included due to inconsistencies found where one ModelSEED ID maps to multiple dissimilar metabolites in another database (e.g., bacterial ubiquinone cpd15560 in ModelSEED maps to both ubiquinone q8 and ubiquinol q8h2 in BiGG; and, similarly, ubiquinol cpd15561 in ModelSEED maps to both metabolites in BiGG). When the algorithm finds a cross-reference pair that corresponds to different universal IDs, the two universal IDs and their properties are merged into a single entity. To avoid cross-reference mapping conflicts, an ID of the source database will not be merged to an ID of the target database that is already mapped to the source database. In this way, priority is given to the first database where the cross-reference was found—hence, the order of the input databases establishes ID precedence for cross-reference conflicts. The algorithm finally outputs the built metabolite and reaction mapper linking the IDs found in any of the databases processed to their universal ID and combined properties. Metabolite and reaction IDs mapping to the same universal ID are considered synonymous. In this way, the universal IDs are used as an internal common namespace to efficiently translate metabolite IDs from different databases and hence enable merging metabolite and reactions across GEMs from different pipelines.

Source databases are updated regularly, and the most recent run of the mapping algorithm processed a total of 2,536,884 metabolite and 299,075 reaction IDs, unifying them into 1,310,929 metabolite and 75,125 reaction unique universal IDs. Figure 2 shows the resultant number of cross-referenced IDs that were mapped by the algorithm (see Supplementary Table). Each of these IDs can thus be recognized by *mergem* and is associated with a unified set of metabolite or reaction properties, including name, InChI key, molecular weight, E.C. number, and chemical formula. The universal mappings and properties can be retrieved and used by third-party applications through the *mergem* Python library.

**Figure 2.**
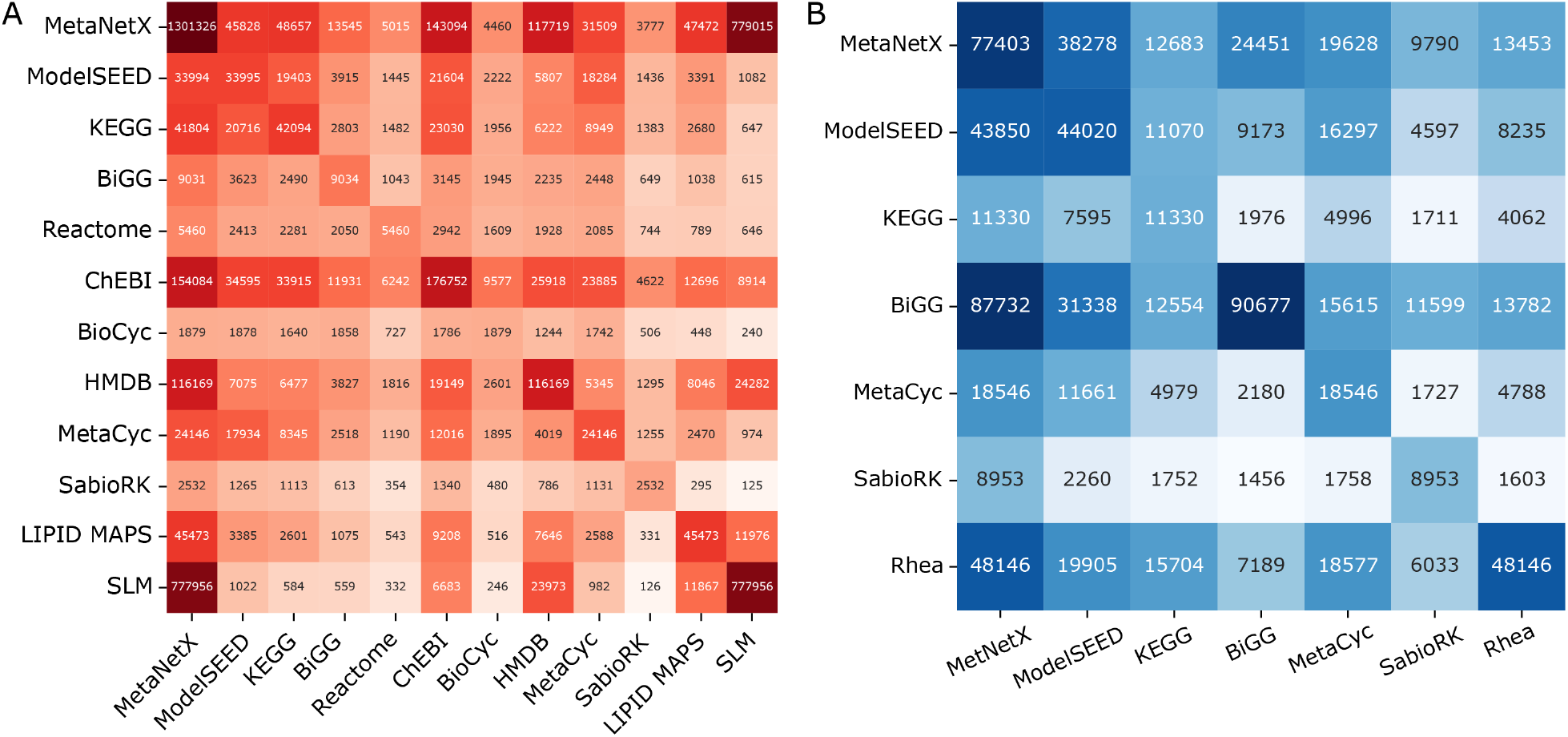
Total number of synonymous metabolite (A) and reaction (B) IDs cross-referenced between databases by the universal mapping system in mergem.

### 3.2. Overview of the merging algorithm

Based on *mergem* universal ID mapper for metabolites (see previous section), we developed an algorithm to merge the metabolites and reactions of two or more GEMs that use the same or different ID namespaces. In brief, the algorithm uses the first model as a template to which the metabolites and reactions from other models are merged into. Metabolite IDs are mapped to a common namespace (called *mergem* ID) as the concatenation of their universal ID and cellular localization. Reactions with the same sets of *mergem* IDs for their reactants and products are considered equivalent and merged among the models. Multiple metabolite IDs can map to the same *mergem* ID, for which the algorithm maximizes metabolite and reaction matches while guaranteeing that all metabolites and reactions from the first model—the template—are present intact in the final merged model (see methods section for details and pseudocode). The objective function for the merged model can be set either by copying the objective function from one of the input models or by merging the objective functions from all the input models.

Once all the input models have been merged, the metabolite IDs are reverted to the original ID from the first model containing it, hence preserving the original ID namespace. The algorithm returns the merged model along with source dictionaries mapping the reactions and metabolites from the input models to the merged model, which can be used for analyzing the commonalities and differences between models. Additionally, a matrix of Jaccard distances is computed and returned by the algorithm (see methods section) that quantifies the similarity between each pair of input models in terms of metabolites (matrix elements above the diagonal) and reactions (matrix elements below the diagonal). The range of each Jaccard distance element is [0,1], where 0 indicates all metabolites or reactions are equal between the models and 1 indicates no common metabolite or reaction exists between the models.

### 3.3. *mergem* package and web application

The presented method is implemented as an open-source Python package called *mergem*. It can be imported into Python scripts or executed as a command-line tool. Models can be input and output in a variety of formats, including SBML (31), MATLAB (The Mathworks, Inc.), and JSON. The efficient implementation of the algorithm can merge standard GEMs in seconds and multiple large models in minutes using a regular desktop computer.

In addition to the Python package, *mergem* has been incorporated into the freely-available web application Fluxer (18), which can create, visualize, and simulate flux networks from GEMs. Fluxer simulates input models with Flux Balance Analysis (FBA) (20) and uses the resulting flux values to compute flux networks that can be visualized as spanning trees, *k*-shortest paths, and complete graphs with multiple layouts. The Fluxer graphs are interactive and the user-friendly interface also allows users to explore large metabolic networks and perform simulations of reaction knock-outs. With the integration of *mergem,* the web-application now allows any user to visually merge and compare any number of GEMs from those already curated in the application—including the models from the BiGG database (32) or previously created in Fluxer through their private URL—or directly from SBML files that can be easily uploaded to the application. Similar to the *mergem* Python package, users can input any number of models for merging, and either select an objective function from any of the input models or merge all objective functions into a single objective reaction.

Figure 3 displays the Fluxer interface after merging three human models: RECON1 (33), red blood cell (iAB_RBC_283 (34)) and platelets (iAT_PLT_636 (35)). The information card on the left includes merging statistics, such as the number of metabolites and reactions merged between any of the input models and the Jaccard distance matrix. The elements below the matrix diagonal (blue) represent the Jaccard distance between the reactions for each pair of models, while the elements above the matrix diagonal (red) represent the Jaccard distance between the metabolites for each pair of models. The diagonal elements are always zero by definition and hence not shown. The intensity of the element colors in the matrix indicates how similar (lighter) or different (darker) the reaction or metabolite sets are between the pair of models. The Fluxer graphs display metabolites, reactions, and their connections in different colors depending on whether they are unique to a particular model (red, purple, and blue in the figure), common to all models (green), or contained in two or more models but not all (gray). Colors can be customized by the user. Clicking on a metabolite or reaction displays a card on the right showing its properties, such as names, molecular weight, links to other databases, and molecular structure, as well as which input models contain that particular reaction or metabolite. Fluxer can simulate the knock-out of reactions in the merged model, after which a new FBA analysis is performed and a new flux graph is computed and displayed. Flux graphs of the merged model can be downloaded as images, vector graphics, and GraphML, and the merged model can be downloaded as SBML. The interface also includes the ability to perform and download a FROG analysis (23), which includes flux variability analysis, reaction deletion fluxes, objective function values, and gene deletion fluxes. Importantly, the URL generated for a merged model is unique and persistent, which can be used to share publicly or privately the merged model including all the interactive functionality and analysis tools in Fluxer.

**Figure 3.**
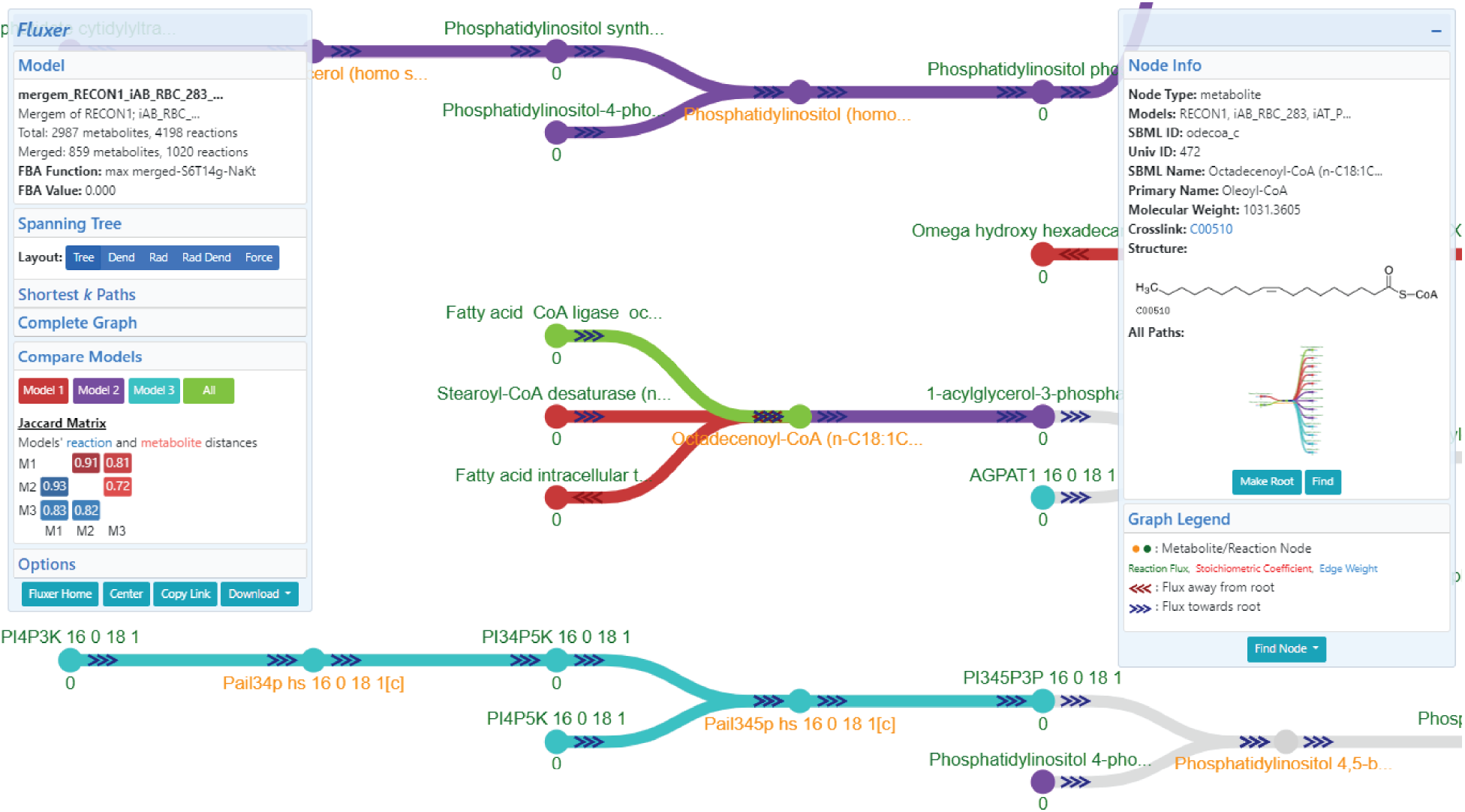
Fluxer interactive web interface after merging three eukaryotic models. Nodes and edges in red, purple, and cyan are unique to RECON1, red blood cells, and platelets, respectively; elements in green are common to all models; and elements in gray belong to only two models. The left card shows details of the merged models and the Jaccard matrix, together with buttons to modify and customize the computed graph. The right card displays properties about the selected metabolite or reaction . Buttons to root, find, or knock-out the selected node are also available in the right card.

### 3.4. Features and performance of *mergem* compared to other tools

Existent tools for merging or comparing GEMs differ in their methodology, their type of interface, the number of file inputs and formats, and their ability to visually display the results. Table 2 compares the main features of the currently available automatic merging tools for GEMs. The main differences are that most tools can merge only pairs of models, are not able to merge models that utilize different ID systems for metabolites and reactions and cannot visually present or allow interaction with the results. MetaMerge (10) was excluded from this study since it is scripted in an outdated version of Python and the package is missing files. ModelBorgifier (11) was also omitted since the method requires the user to manually reconcile metabolites and reactions.

**Table 2.** Key features of currently-available automatic tools for merging and comparing GEMs.

| Tool | Merging method |  | Type | Max. no. models | Model input format | Visualization |
| --- | --- | --- | --- | --- | --- | --- |
|  | Metabolites | Reactions |  |  |  |  |
| **COBRApy (20)** | Direct ID string matching | Direct ID string matching | Python library | 2 | COBRApy model objects | - |
| **MetaNetX (25)** | Internal ID mapper | ID based | Web app | 2 | SBML files following specific standard | - |
| **iMET (12)** | ID based – connects to KEGG for more information before merging | ID based – connects to KEGG for more information before merging | GUI native app | 2 | SBML files | - |
| ***mergem***<br>(This study) | Universal ID mapper | Universal metabolite IDs and their presence as product or reactant | Python command line | Any | SBML, MATLAB, JSON files | - |
|  |  |  | Python library | Any | SBML, MATLAB, JSON files, and COBRApy model objects | - |
| **Fluxer**<br>(This study) | *mergem* algorithm | *mergem* algorithm | Web app | Any | SBML files, select from curated models, URLs to fluxer models | Interactive color-coded visualization |

COBRApy (20) is a Python implementation of COBRA and includes a merging tool that can take as input two COBRA model objects and output a combined model. The merging function performs a direct string matching of the metabolites and reactions IDs to compare pairs of models, which can lead to disconnected networks, duplicated metabolites and reactions, or erroneous deletion of elements in the resulting model. The *mergem* algorithm instead uses a universal mapping dictionary that can translate different database IDs for metabolites to a common namespace. By not using reaction IDs and instead comparing the participating reactant and product metabolites to match reactions, *mergem* bypasses completely the ID mapping problem for reactions which makes it more efficient than COBRApy at merging models.

iMET is a standalone graphical user interface that can semi-automatically merge metabolic networks in the SBML file format (12). The algorithm uses features such as metabolite name and KEGG ID to compare pairs of metabolites after which it assigns a similarity score to each pair. Reactions pairs are also assigned a similarity score by comparing their features, such as reactants and products. The similarity scores are used to reconcile the metabolites and reactions between the two models and users can choose to keep the result for each entity or manually change the merging. This method however can only merge two models at a time, requires the two models to follow the same SBML version, and heavily depends on the information provided within the SBML file itself or information extracted from KEGG (if users select this option) and thus can take hours to finish merging a pair of models. *Mergem* does not have those restrictions or dependencies and takes a few minutes to merge very large models.

MetaNetX is a website with a repository of GEMs that also provides tools to construct, compare, analyze, and simulate GEMs (13). There is no single tool on MetaNetX that helps identify both the common and unique reactions and metabolites between models. Importing each model individually and performing such comparative analyses involves multiple steps on MetaNetX. Further, only two models can be merged at a time on MetaNetX. *Mergem* however can perform simultaneously in a single step the merging and statistical analysis for any number of models.

To evaluate their performance, we used each tool for pairwise-merging six draft models of *Pseudomonas putida* using four different identifier namespaces, as they were reconstructed with different pipelines: AuReMe (36), Pathway Tools (37), CarveMe (38), RAVEN (39), ModelSEED (24), and MetaDraft (40) as provided in Mendoza et al. (19). Figure 4 summarizes the metabolite and reaction merging results for each tool and pair of models and their identifier namespaces (see Supplementary Table 2 for values). Each bar color represents a different tool, while empty circles indicate that the tool failed to complete the merging or loading of one of the input models. The color highlighting the model label indicates the identifier namespace of the model. These results show that *mergem* is not only able to merge models from various reconstruction pipelines using different ID naming systems but also merge most metabolic components. In contrast, iMET and MetaNetX were able to merge only four or three model pairs, respectively. Only the AuReMe and Pathway Tools reconstructions could be merged with all four tools. Visualizing the resulting merged models revealed disconnection in the metabolic networks merged using COBRApy and iMET. Further, the merged models output from iMET use their own IDs and do not follow any database standard. *mergem* thus outperformed all the other tools based on the metabolites and reactions merged, IDs in the merged model, and the extent of integration of the two input models. These results illustrate the importance of using a method more complex than directly matching only reaction IDs and how *mergem* can overcome this problem by using a universal mapper and metabolite-focused approach to match metabolites and reactions.

**Figure 4.**
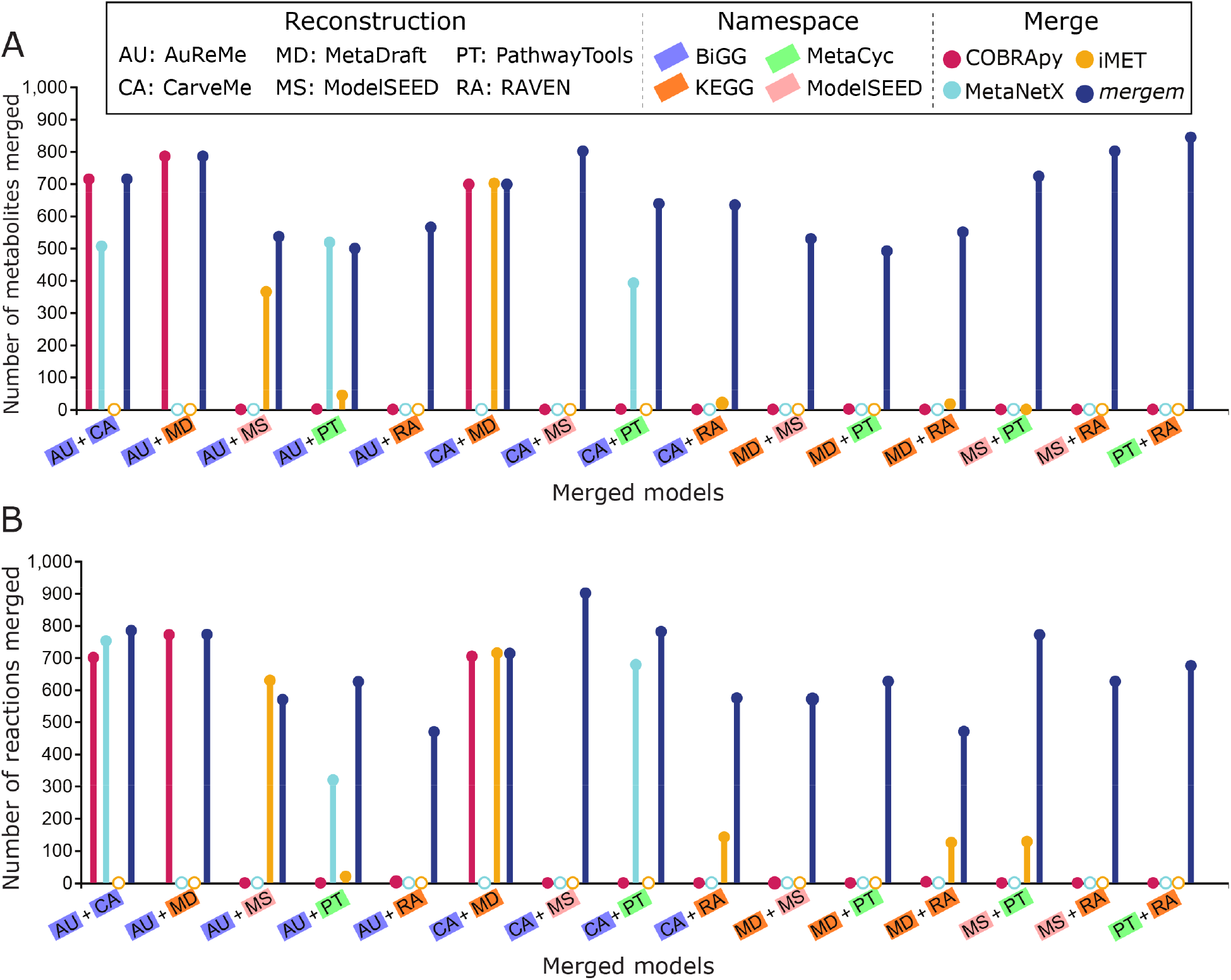
Number of metabolites (A) and reactions (B) merged between pairs of six P. putida reconstructions using four identifier namespaces with four merging tools. Filled circles represent successful merging while empty circles indicate failure to load or merge the models.

### 3.5. Visually comparing GEMs with *mergem* and Fluxer

Differences between metabolic models can arise not only from the use of different reconstruction pipelines but also from different algorithms used for refinement, as well as from models that represent different biological systems. Comparing such models can help study which metabolites and reactions are unique to each model and common between models. The *mergem* algorithm keeps track of the presence of each metabolite and reaction in each input model. Crucially, this information can aid in the investigation of metabolic differences between the cells or organisms that each model represents, which subsequently could be validated experimentally (41), as well as in the evaluation of the metabolic coverage in each reconstruction (42). A graphical visualization of the underlying metabolic networks can further aid in the differential analysis of GEMs.

Fluxer web interface simulates and generates interactive graphs for a given GEM and now it has been extended with the ability to use *mergem* for merging, visually comparing, and analyzing multiple GEMs. The flux network for a merged model can be viewed in the interactive interface as a spanning tree, dendrogram, or complete graph using different layouts. To illustrate this approach for gaining insights regarding metabolic differences, Figure 5 shows the resultant model from merging a platelet (iAT_PLT_636 (35)) and a red blood cell (iAB_RBC_283 (34)) GEM with *mergem* in Fluxer. The nodes and edges in red and blue are unique to the platelet and red blood cell models, respectively. The visualization clearly highlights the different pathways between the two models, such as in the sphingosine and ethanolamine metabolism.

**Figure 5.**
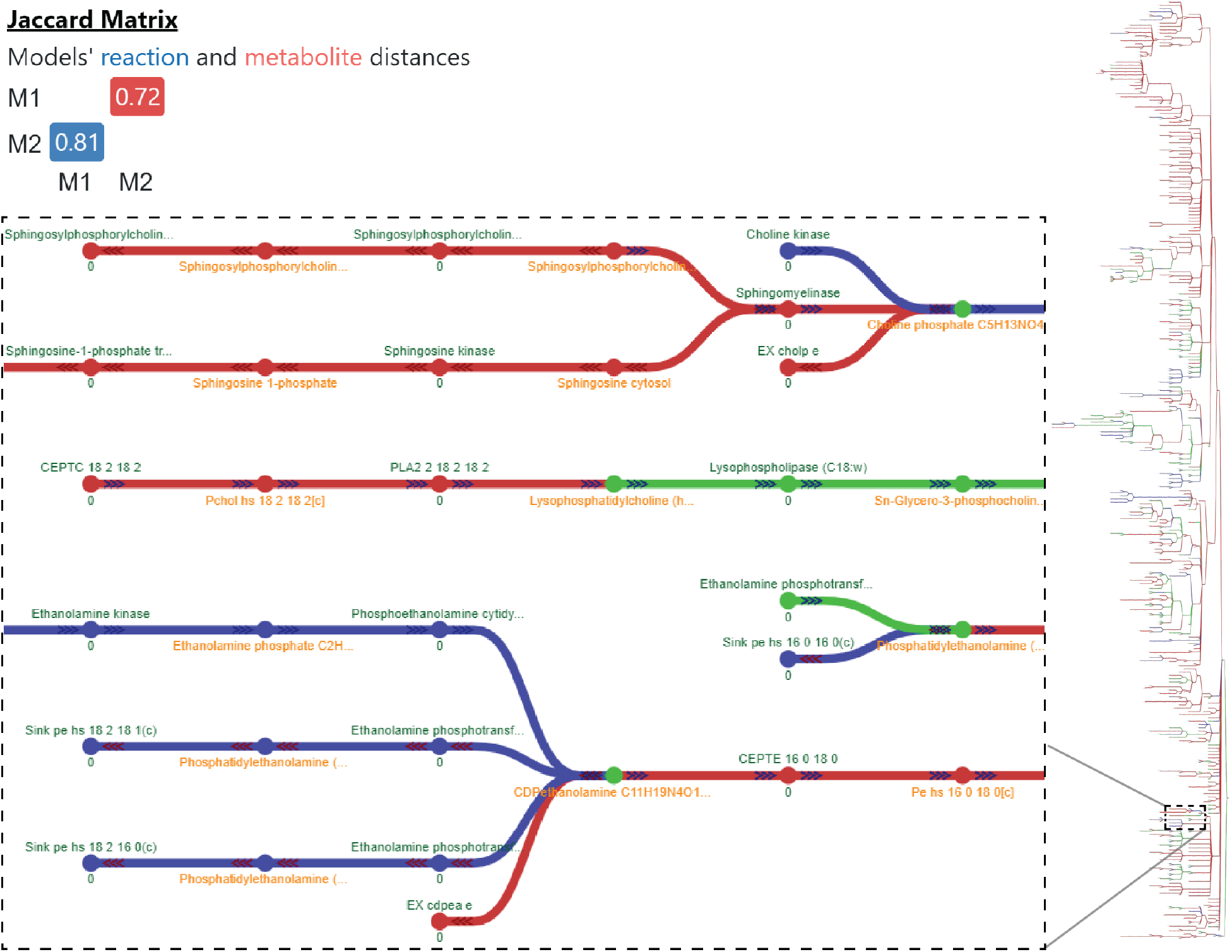
Visual comparison of GEMs for platelets iAT_PLT_636 and red blood cells iAB_RBC_283 with mergem in Fluxer. The results reveal 72% metabolic and 81% reaction differences between the two models, as shown in the Jaccard matrix. The reaction differences include sphingosine and ethanolamine pathways (inset graph). Nodes and edges in blue are unique to red blood cells while those in red are unique to platelet cells. Elements in green are common to both cell types.

### 3.6. Comparing reconstructions using *mergem*

Draft models can vary depending on the pipeline and specific settings used during their reconstructions. Identifying the metabolite and reaction components unique to differently generated reconstructions can aid in curating which components to keep in the final model. There is thus a need for comparing the differences between reconstructions from different pipelines and the integration of the different draft models into a single curated final model. Here we illustrate how *mergem* and Fluxer visualizations can be applied to highlight common and unique elements between draft models.

At the first stage for curating a new GEM, the contents of a draft reconstruction rely on multiple parameters and input data such as the genome sequence, the template model, or gap-filling media. To illustrate how the selection of a different template for a reconstruction can affect the metabolites and reactions added by a pipeline, we compared two *Pseudomonas putida* reconstructions drafted with ModelSEED. The models were generated using either a gram-negative template (MS1) or a core template (MS2) (19). Figure 6 shows the resulting metabolic spanning tree, without zero-flux reactions and cofactor metabolite nodes, after merging and comparing the two draft models with *mergem* in Fluxer. *mergem* found and merged 1730 metabolites (97.5%) and 1666 reactions (95%) in common between the two reconstructions. The analysis revealed that only the model using MS1 contains fatty acid reactions (Figure 6 inset, red nodes) downstream of the acyl carrier protein. These reactions appear to be associated with the metabolism of lipid poysaccharides that are characteristic of gram-negative bacteria for the synthesis of the lipid bilayer (43). The components from gram-negative template are thus evident when comparing the two reconstructions that use different templates.

**Figure 6.**
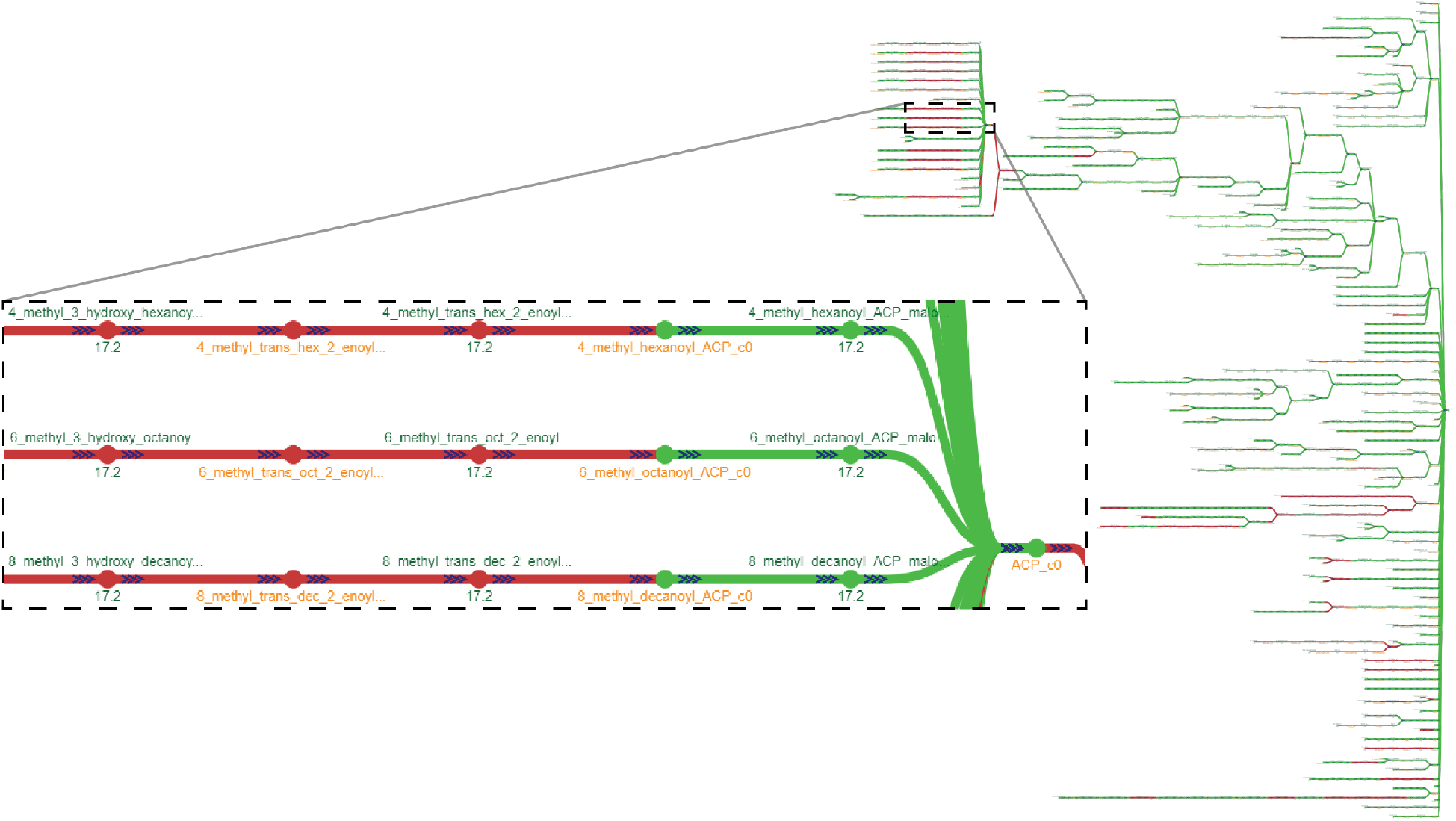
Comparing the effect of reconstruction parameters with mergem in Fluxer. Two different P. putida draft reconstructions built with ModelSEED using either a gram-negative (MS1) or core (MS2) template on ModelSEED reveals components unique to gram-negative template-based models. Flux graph of the complete model visualized as a spanning tree. Green nodes and links indicate metabolic components common to both reconstructions and those in red are unique to the gram-negative template reconstruction. Inset shows some of the unique metabolites and reactions involved in the lipopolysaccharide metabolic pathway characteristic of gram-negative bacteria.

The role of template selection in the CarveMe pipeline was similarly assessed using *mergem* in Fluxer. Two *Lactobacillus plantarum* reconstructions using either a universal bacterial template (CA1) or a gram-positive template (CA2) (19) were merged with *mergem*, resulting in 1031 metabolites (90%) and 1467 reactions (87%) in common between the two reconstructions. Figure 7 shows the resulting metabolic spanning tree, without zero-flux reactions and cofactor metabolite nodes, after merging and comparing the two draft models with Fluxer web interface. The comparison demonstrated that the model from the gram-positive template contained more fatty acid ligase reactions than the one from the universal bacterial template. Additionally, the biomass precursors for the two modes were highly similar (53/61 metabolites in common) except for teichoic acid metabolites, which were unique to the model from the gram-positive template. Indeed, teichoic acid polymers are commonly found in the cell wall of gram-positive bacteria (44, 45). These results demonstrate the importance of template selection when building draft reconstructions and how visually comparing metabolic networks with *mergem* in Fluxer can highlight the components that were added from each reconstruction and where they fit within the overall metabolic flux network.

**Figure 7.**
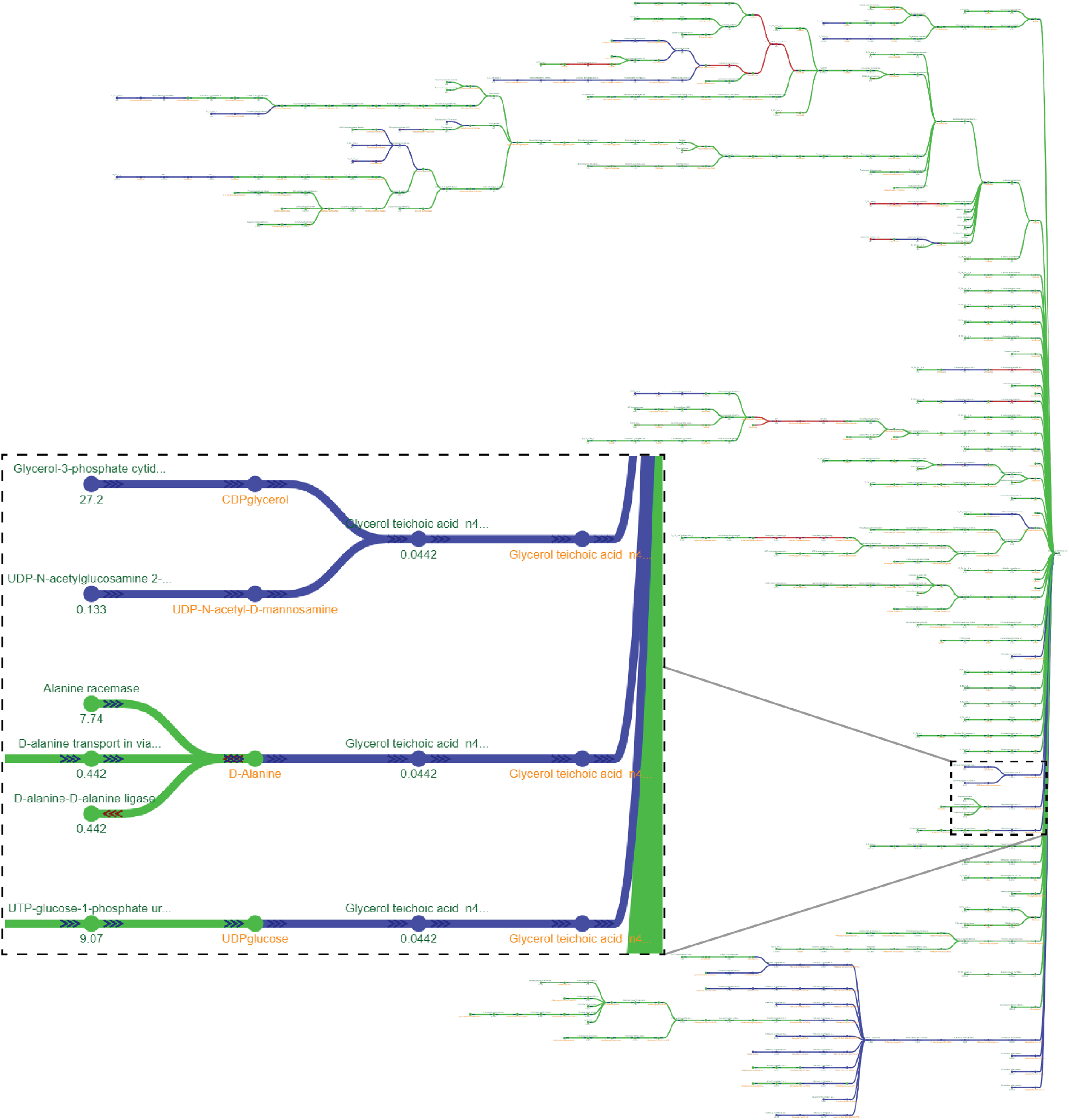
Comparing reconstructions using different templates with mergem in Fluxer. Comparing L. plantarum draft reconstructions using a universal (CA1) or gram-positive (CA2) template in CarveMe reveals specific components unique to gram-positive template-based models. Green nodes and links are metabolic components common to both reconstructions and those in red or blue are unique to the universal or gram-positive models, respectively. Blue components include many techoic acids (insert) that is characteristic of gram-positive bacteria.

### 3.7. Finding missing and newly added reactions using *mergem*

Due to incomplete genome annotations—or errors in them—automatically-generated reconstructions can miss essential reactions and hence fail in producing biomass (indicating growth) when FBA-simulated with an appropriate media (46). A number of tools and algorithms have been developed for finding metabolic and pathway gaps in a reconstruction in base of a set of validated reactions from other organisms. These gap-filling tools include GapFill (47), gapseq (48), and OptFill (49), which Fluxer and *mergem* can complement by visually identifying missing reactions for refining the metabolic model.

Towards the identification of metabolic gaps and possible filling candidates, a draft reconstruction can be compared to a curated model of a closely related organism. To illustrate this approach using *mergem* and Fluxer, a ModelSEED reconstruction for *Lactobacillus plantarum* (19) was merged and compared with a GEM of *Lactobacillus reuteri* (50). Figure 8A shows the results of merging the two models with *mergem* in Fluxer. The ModelSEED reconstruction shared 409 (25%) metabolites and 436 (26%) reactions with the *L. reuteri* model. Using the Fluxer complete graph, with zero-flux reactions and cofactor metabolites hidden, we visually identified nine reactions possibly missing in the ModelSEED reconstruction (Supplementary Table 3). To validate if any of these reactions represent true gaps and good gap-filling candidates, we compared the ModelSEED reconstruction with a new curated model of *L. plantarum* (19). Figure 8B shows the results, revealing that all nine reactions were required to metabolically complete the ModeSEED reconstruction. Inset graphs in Figure 8A and Figure 8B show one of the potential gaps and fillers identified in the analysis. While not all metabolic gaps need to be filled in a final model, *mergem* and Fluxer visualizations represent a crucial aid to improve and assess the completeness of models by comparing them to more recent curations of closely related organisms.

**Figure 8.**
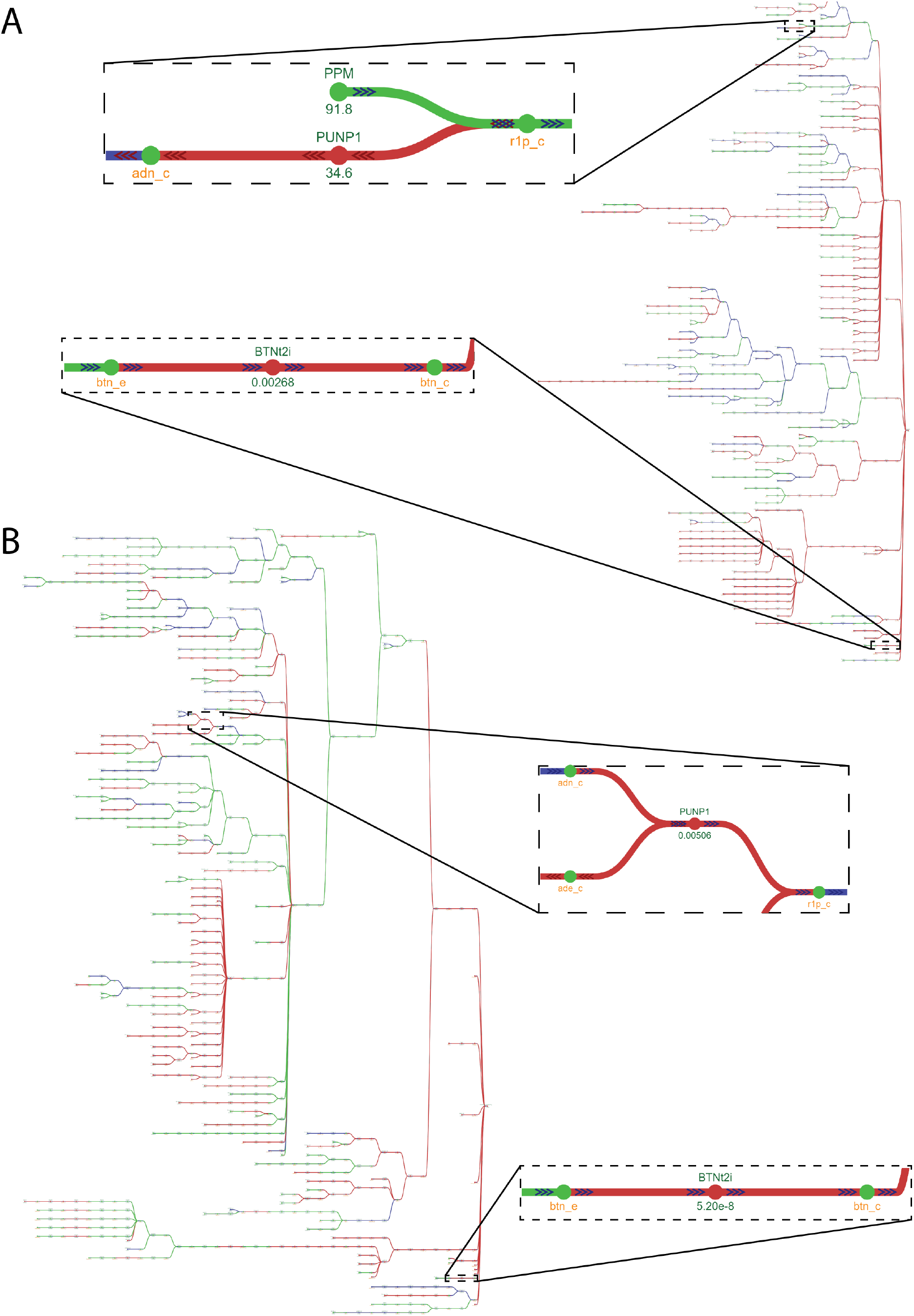
mergem and Fluxer can aid in the identification of possible gaps and filling candidates in draft reconstructions. **A.** Comparing a ModelSEED draft reconstruction for Lactobacillus plantarum with a manually curated model for Lactobacillus reuteri revealed possible metabolic gaps in the draft model and potential reactions that can fill them. Nodes and links in red are unique to the manually-curated L. reuteri model, while those in green are common between the two models. **B.** Comparing the same ModelSEED draft reconstruction for L. plantarum with a manually curated model for the organism showed the same reactions as missing in the reconstruction, suggesting that the proposed gaps and fills were correct. The nodes and links in red are unique to the manually curated model and those in green are common between the draft and curated models. Paths zoomed in the inset boxes contain examples of gaps and its filling candidates (Purine nucleoside phosphorylase, PUNP1 and Biotin exchange, BTNt2i) found with this approach.

Tools and algorithms for gap-filling and other refinement methods most often output the updated models without explicitly listing the newly added reactions and metabolites. In addition, published GEMs are regularly updated when more information is available regarding the corresponding biological system. Comparing a model that has undergone refinement with its previous version can provide insights regarding the metabolic elements that were added as part of the refinement process. To demonstrate how *mergem* and Fluxer can be used to identify updates on models, even when they use different ID namespaces, we downloaded, merged, and compared three models of *Pseudomonas putida* KT2440: iJN1463, the most recently published model for this strain, which is a refinement adding and deleting reactions and metabolites across multiple *P. putida* models (51); iJN746, the first published GEM for this strain (52); and MNX_iJN746, a translation of iJN746 to the MetaNetX namespace as retrieved from MetaNetX webserver (13). Figure 9A shows the resultant comparison by *mergem* with a flux graph in Fluxer, together with its computed Jaccard matrix. The results show how the original model and the version published in MetaNetX, based on a different ID namespace, are highly similar (99% metabolite match and 99.8% reaction match), but they both differ in 60% and 68% of metabolites and reactions, respectively, with respect to the refined model. This shows how *mergem* can successfully merge and compare models in different database namespaces and highlight differences between model refinements. Aided by the visual comparison in Fluxer, we easily identified that the elements added to the refined model included fatty acids pathways, pyrroloquinoline quinone synthesis, and iron metabolism, as reported in (51). The inset graph in Figure 9A highlights in red, as visualized in Fluxer, some of the newly-added reactions and metabolites in the refined model. The nodes and links in gray (such as dimethylglycine dehydrogenase reaction, DMGDH) that are common only to the *P. putida* models prior to refinement (both original, iJN746 and its MetaNetX conversion, MNX_iJN746) indicate their deletion or update as part of the model refinement. A closer look at the dimethylglycine dehydrogenase reactions revealed that the older versions of *P. putida* GEMs did not contain nicotinamide dehydrogenase reduction as part of the reaction while the same reaction in the most recent model has been updated to include this cofactor conversion. Similar visual comparisons can be performed with *mergem* for reconstructions before and after gap-filling reactions are applied or for reconstructions obtained from different gap-filling algorithms.

**Figure 9.**
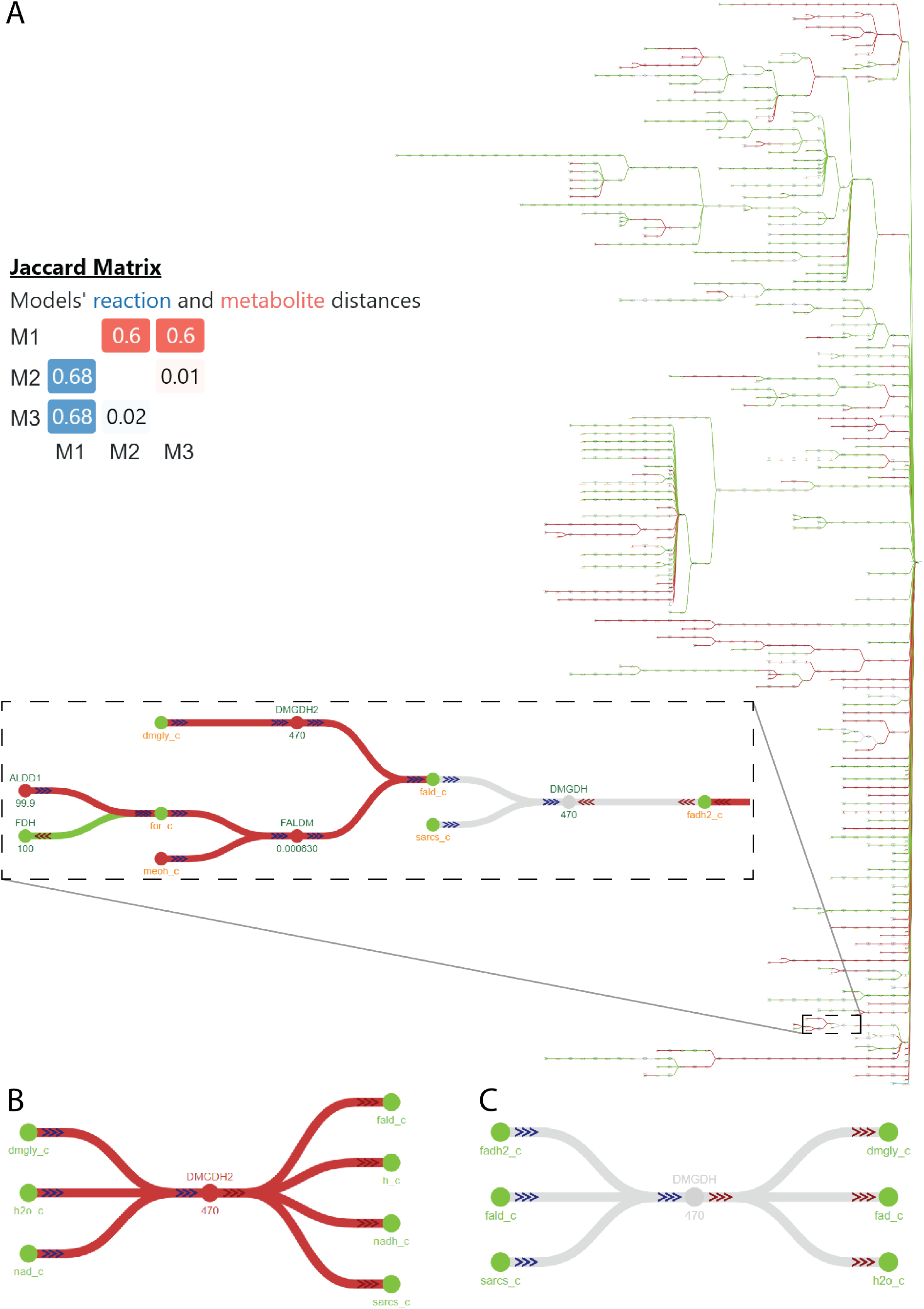
Model update analysis using mergem and Fluxer with input models that follow different database identifier namespaces. Updates performed during refinements for three P. putida KT2400 models: most recent model M1:iJN1463, first published model M2:iJN746 and the first published model in MetaNetX namespace M3:MNX_iJN746. **A.** Spanning tree graph of merged model and its resultant Jaccard matrix. Added, deleted, and retained components are visible in red, gray, and green, respectively. **B.** The reaction dimethylglycine dehydrogenase updated to include the reduction of nicotinamide adenine dinucleotide. **C.** In comparison, the reaction dimethylglycine dehydrogenase before the update.

### 3.8. Comparing closely related organisms using *mergem*

Organisms of different species but same genus often show many similar but a few unique metabolic phenotypes. Using *mergem* with GEMs of such organisms can aid in globally comparing the metabolisms between related species. To illustrate this case, we compared a GEM for *Pseudomonas aeruginosa* iPau21 (53) with two GEM versions of *Pseudomonas putida* (iJN1463 (51) and iJN746 (52)). iJN746 is the first published model for *P. putida, while* iJN1463 is a refinement of iJN746 and thus contains updated reaction and metabolite information for *P. putida.* Figure 10A shows the complete tree graph for the resultant merged model in radial layout as visualized in Fluxer and the Jaccard matrix resulting from *mergem* comparison. As expected, the two models for *P. putida* (iJN1463 and iJN746) were more similar to each other than to the model for a different species (iPau21). The results further show that the *P. aeruginosa* model is more similar to the older *P. putida* model (iJN746) than its refined counterpart (iJN1463). One cause for this could be the lack of species-specific information at the time of construction of iJN746. Of the 116 biomass precursors in the merged model, 67 (57%) were common between all models, 7 (0.06%) were unique to *P. aeruginosa*, and 18 (15%) were unique to the refined *P. putida* model, iJN1463. None of the precursors were unique to the *P. putida* model prior the refinement, iJN746. The biomass precursors unique to *P. aeruginosa* included *ubiquinol-9, protein, RNA, Pseudomonas LPS core*, and *peptidoglycan polymer*. Although *P. putida* and *P. aeruginosa* belong to the same genus, they show differences in certain pathways. For example, *P. aeruginosa* contains pathways for production of virulence factors that include alginate (Figure 10B, red elements), rahmnolipids, and phenezines. In the merged model network, the reactions unique to *P. aeruginosa* were primarily lipid reactions (Figure 10C, red elements). Similarly, *P. putida* contains pathways metabolizing aromatic compounds such as toluene, indole, and m-xylene (Figure 10D, gray and purple elements). Similar analyses can be performed to distinguish metabolic features of different organisms.

**Figure 10.**
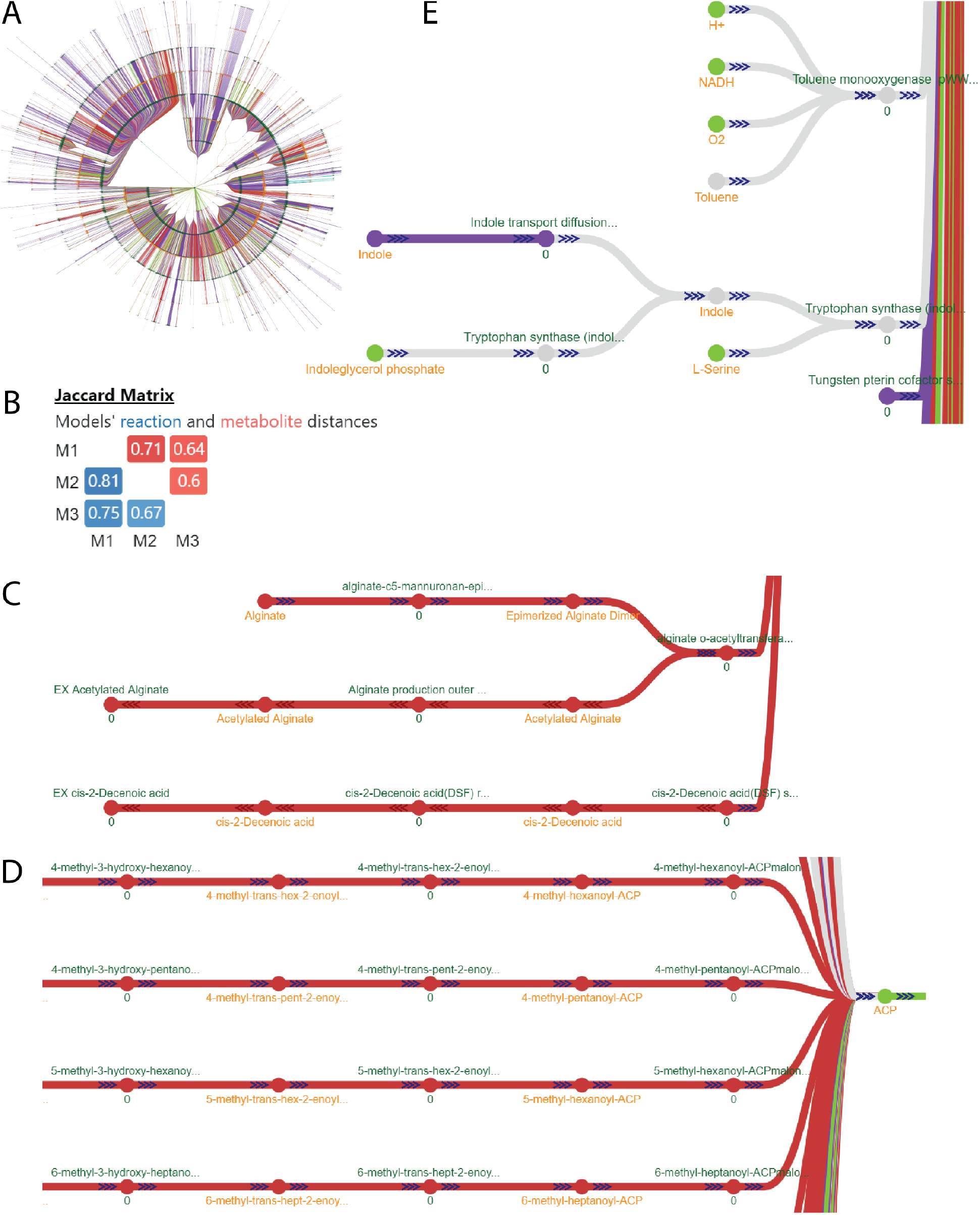
The metabolism of different organisms can be contrasted by merging and comparing their GEMs with mergem in Fluxer. A comparison of P. aeruginosa (model iPau21) with two models of P. putida (iJN1462 and its previous version as iJN746) reveals their common and unique pathways. **A.** Complete graph of the merged model in radial layout and the resulting Jaccard distance matrix (M1: iPau21, M2: iJN1462, and M3: iJN746). **B.** Alginate pathway reactions found to be unique to P. aeruginosa. **C.** Some of the fatty acid pathway reactions found to be unique to P. aeruginosa. **D.** Toluene and indole and their associated reactions in gray or purple are present only in the P. putida models. Elements in green are common to all models; elements in red are unique to P. aeruginosa; elements in purple are unique to iJN1462; elements in gray are common to two models.

### 3.9. Comparing organisms for finding common drug targets

Polymicrobial infections can be more severe and difficult to treat than infections of the same species alone (54). GEMs can facilitate the discovery of potential antimicrobial drug targets (55), and comparing GEMs of pathogens to find commonalities can aid in the identification of shared antibacterial targets as candidates for effective treatments in co-infections. To illustrate this application, we used *mergem* in Fluxer to visualize the commonalities between the curated GEMs for three gram-negative pathogens: *Acinetobacter baumanii* (56)*, Klebsiella pneumoniae* (57), and *Pseudomonas aeruginosa* (53). Figure 11 shows the visualization of the resultant merged models in Fluxer. Since the three gram-negative pathogens contained around 60% unique reactions and 70% unique metabolites, as shown in the Jaccard matrix (Figure 11), we hypothesized that pathways common to all three pathogens could be candidates for antimicrobial drugs. Among the common elements, this comparison revealed the shikimate and riboflavin pathways (Figure 11, green), which are indeed targets of current antimicrobials (58, 59). The other reactions found in common are good candidates for further study to test their effectiveness as antibacterial targets against gram-negative bacterial pathogens.

**Figure 11.**
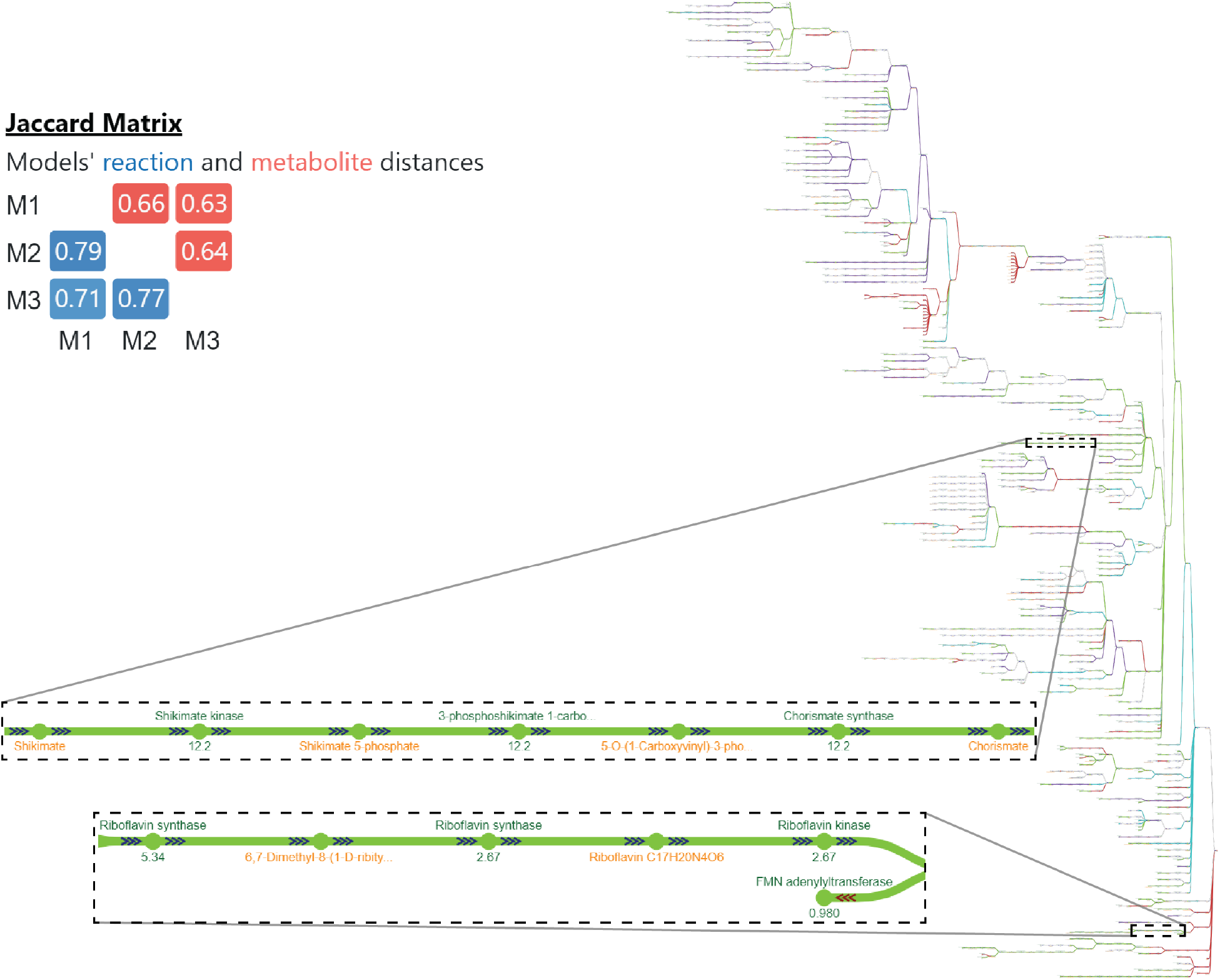
Potential antibacterial targets were identified with mergem in Fluxer by comparing GEMs of three gram-negative pathogens, including Acinetobacter baumanii, Klebsiella pneumoniae, and Pseudomonas aeruginosa. The resulting common pathways included shikimate and riboflavin (insets), which hence are good candidates as antimicrobial targets.

## 4. Discussion

Here we presented *mergem*, a novel tool for merging and comparing GEMs. The method computes and uses a universal database ID mapping dictionary to translate metabolite IDs from multiple different database systems to a common namespace. This allows reactions to be merged by comparing the translated metabolite IDs of their reactants and products. *mergem* can be used on the command-line or imported into Python scripts. While there exist other tools that can merge GEMs, *mergem* shows improvement over these methods in terms of recognition and reconciliation of metabolite IDs, merging of reactions based on these metabolite mappings, and range of model inputs.

To enable the visual and interactive comparison of models and the construction of merged models with a user-friendly tool, we incorporated the presented methodology into the web-application Fluxer. This approach can create, visualize, and simulate interactive flux graphs of multiple merged models in which metabolites and reactions are color-coded depending on their model source. We demonstrated this visual methodology for merging and comparing draft reconstructions from different pipelines, even when the models use different database IDs. In addition, we showed applications of the methodology for visually identifying potential gaps in reconstructions and for finding commonalities useful for discovering antimicrobial targets. The tidy, interactive flux graphs produced by Fluxer with *mergem* allow users to visually inspect the differences between models and, crucially, understand the location of such differences in the context of the whole metabolic flux network of an organism or cell.

Discrepancies between GEMs and reconstructions for the same organism or cellular system can arise from any of the steps during curation: (1) genome annotation, (2) environment specification, (3) biomass formulation, (4) network gap-filling, and (5) flux simulation method (60). Merging and visually comparing the metabolic models at each stage of the reconstruction process using the proposed methodology with *mergem* and Fluxer, could help in the detection of erroneous or missing metabolic data in the models. The examples described illustrate how *mergem* can aid in resolving these five sources of discrepancies, from finding missing reactions by comparing close-related organisms to validating a curated model with a FROG report using different simulation methods. Future work will extend the *mergem* algorithm to include gene expression, metabolomic, and fluxomic data for comparing metabolic networks. Such extensions will further increase our ability for visualizing and contrasting different metabolic phenotypes.

GEMs are essential tools for understanding and predicting metabolic behaviors in research and engineering applications. Making systems-level comparisons of metabolism using GEMs requires models that are comprehensive and accurate in their biological representation. The process of building such genome-scale models involves working with many drafts and refining them iteratively. Unfortunately, different reconstruction tools use different database IDs, which makes it challenging to compare and integrate models from different pipelines. While there is no gold standard pipeline for generating comprehensive drafts, we showed that using *mergem* to integrate drafts from multiple reconstruction methods could be a valuable aid for model curation, including efforts to build universal community models such as the Human Metabolic Atlas (7). While *mergem* represents a significant advancement for accurately mapping IDs from several databases, there is still a need for a global standardization in metabolite and reaction IDs and definitions. Such standardization certainly would improve merging tools such as *mergem* and facilitate the curation of comprehensive GEMs and their comparison.

## 5. Conclusions

*mergem* and its interactive visual integration in Fluxer enable new comparative studies of GEMs for different reconstructions, curated models, and organisms. In addition to providing a means to construct standardized and comprehensive GEMs, the presented tools will be an aid for generating hypotheses regarding the genetic targets that can be overexpressed or knocked-out to optimize any phenotype, such as those in disease-relevant pathways or for the production of value-added metabolites in engineering. Future applications will include using *mergem* and Fluxer to build and analyze robust GEMs models that can predict optimized growth phenotypes to be further integrated with mechanistic models and machine-learning methodologies (61–64).

## 6. Declarations

### Data Availability

*mergem* can be freely installed from PyPI with *pip*, the package installer for Python. The source code is available in GitHub (https://github.com/lobolab/mergem) and Zenodo (https://doi.org/10.5281/zenodo.7865478). The universal mapping system in *mergem* for metabolites and reactions can be exported as CSV files and automatically updated with the latest external database information. Detailed documentation for using *mergem* is available at https://mergem.readthedocs.io and a manual is included as supplementary information. All the models shown in the figures are freely available to visualize, analyze, and download in Fluxer at https://fluxer.umbc.edu/ and their URLs are listed in Supplementary Table 4.

### Authors’ contributions

AH and DL designed and implemented the method and wrote the manuscript. AH analyzed the data. DL secured funding. All authors read and approved the final manuscript.

## Acknowledgements

We thank Daniel Machado for useful discussions, Ivan Erill and Jeffrey Gardner for critical reading of the manuscript, and the members of the Lobo Lab and the students and faculty at UMBC for their support and help in testing the software applications.

