## Supplementary Tables for "*mergem*: merging and comparing genome-scale metabolic models using universal identifiers"

**Contents**


| | |
|---|---|
| Supplementary Table 1 | Number of metabolite and reaction IDs cross-referenced by the universal mapping system. |
| Supplementary Table 2 | Number of metabolites and reactions merged between pairs of reconstructions using different tools. |
| Supplementary Table 3 | Potential metabolic gaps in ModelSEED reconstruction for *L. plantarum* and candidates for filling the gaps. |
| Supplementary Table 4 | Fluxer URLs for each of the models presented in the main figures. |

**Supplementary Table 1.** Number of metabolite and reaction IDs cross-referenced by the universal mapping system. Rows and columns indicate source and target databases, respectively. Diagonal indicates total number of IDs in each database.

| Metabolites | MetaNetX | ModelSEED | KEGG | BiGG | Reactome | ChEBI | BioCyc | HMDB | MetaCyc | SabioRK | LIPID MAPS | SLM |
|---|---|---|---|---|---|---|---|---|---|---|---|---|
| MetaNetX | 1301326 | 45828 | 48657 | 13545 | 5015 | 143094 | 4460 | 117719 | 31509 | 3777 | 47472 | 779015 |
| ModelSEED | 33994 | 33995 | 19403 | 3915 | 1445 | 21604 | 2222 | 5807 | 18284 | 1436 | 3391 | 1082 |
| KEGG | 41804 | 20716 | 42094 | 2803 | 1482 | 23030 | 1956 | 6222 | 8949 | 1383 | 2680 | 647 |
| BiGG | 9031 | 3623 | 2490 | 9034 | 1043 | 3145 | 1945 | 2235 | 2448 | 649 | 1038 | 615 |
| Reactome | 5460 | 2413 | 2281 | 2050 | 5460 | 2942 | 1609 | 1928 | 2085 | 744 | 789 | 646 |
| ChEBI | 154084 | 34595 | 33915 | 11931 | 6242 | 176752 | 9577 | 25918 | 23885 | 4622 | 12696 | 8914 |
| BioCyc | 1879 | 1878 | 1640 | 1858 | 727 | 1786 | 1879 | 1244 | 1742 | 506 | 448 | 240 |
| HMDB | 116169 | 7075 | 6477 | 3827 | 1816 | 19149 | 2601 | 116169 | 5345 | 1295 | 8046 | 24282 |
| MetaCyc | 24146 | 17934 | 8345 | 2518 | 1190 | 12016 | 1895 | 4019 | 24146 | 1255 | 2470 | 974 |
| SabioRK | 2532 | 1265 | 1113 | 613 | 354 | 1340 | 480 | 786 | 1131 | 2532 | 295 | 125 |
| LIPID MAPS | 45473 | 3385 | 2601 | 1075 | 543 | 9208 | 516 | 7646 | 2588 | 331 | 45473 | 11976 |
| SLM | 777956 | 1022 | 584 | 559 | 332 | 6683 | 246 | 23973 | 982 | 126 | 11867 | 777956 |

| Reactions | MetaNetX | ModelSEED | KEGG | BiGG | MetaCyc | SabioRK | Rhea |
|---|---|---|---|---|---|---|---|
| MetaNetX | 77403 | 38278 | 12683 | 24451 | 19628 | 9790 | 13453 |
| ModelSEED | 43850 | 44020 | 11070 | 9173 | 16297 | 4597 | 8235 |
| KEGG | 11330 | 7595 | 11330 | 1976 | 4996 | 1711 | 4062 |
| BiGG | 87732 | 31338 | 12554 | 90677 | 15615 | 11599 | 13782 |
| MetaCyc | 18546 | 11661 | 4979 | 2180 | 18546 | 1727 | 4788 |
| SabioRK | 8953 | 2260 | 1752 | 1456 | 1758 | 8953 | 1603 |
| Rhea | 48146 | 19905 | 15704 | 7189 | 18577 | 6033 | 48146 |

**Supplementary Table 2.** Number of metabolites and reactions merged between pairs of models using different tools. "-" indicates tool failure when loading or merging the models. AU: AuReMe, CA: CarveMe, MS: ModelSEED, MD: MetaDraft, PT: Pathway Tools, and RA: RAVEN

| **Metabolites** | **COBRApy** | **MetaNetX** | **iMET** | ***mergem*** |
|---|---|---|---|---|
| AU + CA | 714 | 506 | - | 714 |
| AU + MD | 785 | - | - | 785 |
| AU + MS | 0 | - | 365 | 536 |
| AU + PT | 1 | 518 | 44 | 499 |
| AU + RA | 0 | - | - | 565 |
| CA + MD | 698 | - | 701 | 698 |
| CA + MS | 0 | - | - | 801 |
| CA + PT | 1 | 392 | - | 638 |
| CA + RA | 0 | - | 19 | 634 |
| MD + MS | 0 | - | - | 529 |
| MD + PT | 1 | - | - | 491 |
| MD + RA | 0 | - | 17 | 550 |
| MS + PT | 0 | - | 0 | 723 |
| MS + RA | 0 | - | - | 801 |
| PT + RA | 0 | - | - | 844 |

| **Reactions** | **COBRApy** | **MetaNetX** | **iMET** | ***mergem*** |
|---|---|---|---|---|
| AU + CA | 700 | 752 | - | 784 |
| AU + MD | 771 | - | - | 772 |
| AU + MS | 0 | - | 629 | 569 |
| AU + PT | 0 | 319 | 20 | 625 |
| AU + RA | 3 | - | - | 469 |
| CA + MD | 704 | - | 714 | 713 |
| CA + MS | 0 | - | - | 901 |
| CA + PT | 0 | 678 | - | 781 |
| CA + RA | 0 | - | 142 | 574 |
| MD + MS | 0 | - | - | 571 |
| MD + PT | 0 | - | - | 626 |
| MD + RA | 3 | - | 125 | 470 |
| MS + PT | 0 | - | 128 | 771 |
| MS + RA | 0 | - | - | 626 |
| PT + RA | 0 | - | - | 675 |

**Supplementary Table 3.** Nine potential gaps and fillers identified in ModelSEED reconstruction for *L. plantarum*. Manually curated model for *L. reuteri* from (Kristjansdottir *et al.*, 2019). ModelSEED reconstruction and manually curated *L. plantarum* models from (Mendoza *et al.*, 2019).

| Reaction ID | | Reaction name |
|---|---|---|
| ***L. reuteri*** | ***L. plantarum*** | |
| ABTA | ABTA | 4-Aminobutyrate transaminase |
| GLUT6 | GLUt2r | L-Glutamate transport via proton symport |
| PUNP1 | PUNP1 | Purine nucleoside phosphorylase-Adenosine |
| PUNP3 | PUNP3 | Purine nucleoside phosphorylase-Guanosine |
| PUNP5 | PUNP5 | Purine nucleoside phosphorylase-Inosine |
| ALATA_Lr | ALATA_Lr | Alanine transaminase |
| BTNt2i | BTNt2i | Biotin uptake |
| MALTt | MALTt2 | Maltose transport via proton symport |
| HYPOE | HYPOE | Pyridoxamine-5'-phosphate phosphohydrolase |

**Supplementary Table 4.** URLs for the models presented in each of the main figures, which can be used to access, analyze, and download the models from Fluxer web application.

| Figure | Fluxer URL |
|---|---|
| 3 | https://fluxer.umbc.edu/model?id=1fb050032ca2e1dce96190af995cadb92b40f184_3963a7426d7d662b0f404284c397e13227361d88_0335833002b8337f8440cf1f09a41c9aa6d3268f_obj_merge |
| 5 | https://fluxer.umbc.edu/model?id=0335833002b8337f8440cf1f09a41c9aa6d3268f_3963a7426d7d662b0f404284c397e13227361d88_obj_merge |
| 6 | https://fluxer.umbc.edu/model?id=7fe8a8e65427f5f30412cc3341b5ec596e956f42_8483528fcd5891b944d53a6e4f61214acb596f42_obj_merge |
| 7 | https://fluxer.umbc.edu/model?id=631396a6150696a9af6f26b9bca4c9b63343b99f_f25b134379c57bba2fe256c7ad737f8729cb2864_obj_merge |
| 8A | https://fluxer.umbc.edu/model?id=fb168cce92cd64a84ecbbf4e13dedad56adadedb_bd1605b71563d7dde27f5f57552651d9934c0333_obj_1 |
| 8B | https://fluxer.umbc.edu/model?id=6002faacf35a41f4f1b4132f045dd6ff7071d31e_bd1605b71563d7dde27f5f57552651d9934c0333_obj_1 |
| 9 | https://fluxer.umbc.edu/model?id=f4f30bd4265c5734d98b719ce39e0dbbd5d4ecfb_0bab56f8be08a3d62f24dd16c00b6b01fd85cb27_7fe25baa10e0f3ee212b5b8b4edc4742e5ebfd8b_obj_merge |
| 10 | https://fluxer.umbc.edu/model?id=5f84c87ff0449775359f3adb1ddfc75d6e96202d_f4f30bd4265c5734d98b719ce39e0dbbd5d4ecfb_0bab56f8be08a3d62f24dd16c00b6b01fd85cb27_obj_merge |
| 11 | https://fluxer.umbc.edu/model?id=e49e478a9849adcb1cb7e03409de63e470b2880d_a84d145f63822725400bac076aa22078d6a77189_5f84c87ff0449775359f3adb1ddfc75d6e96202d_obj_1 |
